# On the mechanism of calcium permeability and magnesium block in NMDA receptors - a central molecular paradigm in neuroplasticity

**DOI:** 10.1101/2025.11.06.686637

**Authors:** Ruben Steigerwald, Max Epstein, Tsung-Han Chou, Hiro Furukawa

## Abstract

Neuroplasticity is a fundamental cellular mechanism underlying learning and memory formation and is primed by the coincidental detection of neurotransmitter release from the presynapse and the subsequent calcium influx upon voltage change in the postsynaptic membrane (Bliss and Collingridge, 1993). Molecular assemblies that achieve these events are N-methyl-D-aspartate receptors (NMDARs), which bind the neurotransmitter glutamate and a co-agonist, either glycine or D-serine, and allow Ca^2+^ influx upon relief of the Mg^2+^ channel blockade by membrane depolarization. However, the molecular basis governing Ca^2+^ permeability and Mg^2+^ blockade in NMDAR remains limited. Here, we demonstrate that Ca²⁺ permeation through the narrow constriction of the cation selectivity filter involves partial dehydration, as evidenced by multiple Ca²⁺ binding sites captured using single-particle cryo-electron microscopy (cryo-EM). In contrast, Mg^2+^ binds outside of the selectivity filter through the water network by remaining hydrated, thereby serving as a channel blocker. Furthermore, we show that the lipid network around the selectivity filter influences the stability of Mg^2+^ binding. Our study details the critical transmembrane chemistry of NMDAR for initiating neuroplasticity.

## Introduction

The fundamental electro-cellular event initiating the classic Hebbian plasticity for learning and memory is the coincidental detection of neurotransmitters and a voltage change in the postsynaptic membrane(Bliss and Collingridge, 1993; Malenka and Bear, 2004). The postsynaptic NMDARs are pivotally involved in this process by facilitating high calcium influx upon glutamate binding(MacDermott et al., 1986) when the channel blockade by magnesium is relieved by depolarization(Mayer et al., 1984; Nowak et al., 1984), thereby triggering downstream signaling only when the presynaptic and postsynaptic neurons activate concurrently. However, the structural mechanism by which NMDAR channels distinguish calcium and magnesium ions remains unanswered. NMDARs are heterotetrameric receptors comprised of two glycine-binding GluN1 subunits and two glutamate-binding GluN2 subunits (GluN2A-D)(Hansen et al., 2021). The NMDAR channel activation requires multiple layers of conditions, including the occupancy of GluN1 ligand-binding domain (LBD) with a co-agonist, glycine or D-serine, glutamate binding to the GluN2 LBD upon neurotransmission, and depolarization of the postsynaptic membrane to relieve the Mg^2+^ channel block at the transmembrane domain (TMD). The opening of the NMDAR channels results in the conductance of Na^+^, K^+^, and, importantly, Ca^2+^, triggering the subsequent neuroplastic signaling via CaMKII activation (Lisman et al., 1997; Yasuda et al., 2022). The spatial- temporal regulation of the Ca^2+^ influx in neurons controls various forms of plasticity (Jain et al., 2024; Yasuda et al., 2022). Therefore, Ca^2+^ permeation and voltage-dependent Mg^2+^ block in NMDAR are primers for neuroplasticity in excitatory neurons. Despite advances in the structural biology of NMDARs over the past decade (Karakas and Furukawa, 2014; Lee et al., 2014; Zhou and Tajima, 2023), understanding how the NMDAR channel pore distinguishes between Ca^2+^ and Mg^2+^, two structurally similar divalent cations, has remained a challenge. Here, we resolve this classic principle through high-resolution structural biology, showing that Ca^2+^ has at least five binding sites within the cation selectivity filter, likely involving partial dehydration. In contrast, there are strictly two binding sites for hydrated Mg^2+^, at the extracellular and intracellular entrances of the selectivity filter.

### Cryo-EM captures the multiple calcium sites within the NMDAR channel pore

We first sought to capture the calcium permeation pathway within the NMDAR by resolving the structure of the GluN1a-2B NMDAR bound to agonists (glycine and glutamate) in the presence of CaCl_2_ (10 mM) and in the absence of a divalent cation (1 mM EDTA). We selected the GluN1a-2B NMDAR subtype, as it exhibits the highest Ca^2+^ permeability and sensitivity to Mg^2+^ block, similar to the GluN1a-2A NMDAR. The GluN1a-2C and GluN1a-2D NMDARs are significantly less sensitive to Ca^2+^ and Mg^2+^ (Kuner et al., 1996; Siegler Retchless et al., 2012).

Our single-particle cryo-EM achieved TMD resolution ranging from 2.3 to 3.0 Å by combining extensive 3D classification followed by local refinement of the TMD channel region (**Fig. 1a-b, Extended Data Fig. 1**). This allowed us to monitor Ca^2+^ and tightly bound water oxygens at distinct positions in the pore (**Fig. 1, Extended Data Fig. 1**). The single-particle analysis strategy implemented here (**Extended Data Fig. 1**) was crucial in unveiling the hydration pattern of the NMDAR channel pore and Ca^2+^ at multiple positions (**Fig. 1d-e**). Our analysis revealed Ca^2+^ ions within the narrow constriction formed by the re-entry loops located between the M2 and M3 helices of GluN1a and M2’ and M3’ helices of GluN2B (**Fig. 1b**). This motif contains a cluster of Asn (N) residues and is classically annotated as the QRN site, determining the pattern of calcium permeability in ionotropic glutamate receptors (iGluRs) (**Fig. 1c**) (Burnashev et al., 1992b; Sommer et al., 1991). For example, the equivalent site is Gln (Q) in calcium-permeable (cp) AMPARs, whereas it is Arg (R) in calcium-impermeable AMPARs and kainate receptors. However, a recent study demonstrated that the R-form of GluA2 AMPAR can also permeate Ca^2+^ ions when combined with GluA1 and auxiliary subunits such as TARP gamma-2 (Miguez-Cabello et al., 2025). The N-site in GluN1a-2B NMDAR has a narrow cage-like structure formed by the side chain of GluN1a Asn616 and main and side chains of GluN2B Asn615-616 (**Asn-cage; Fig. 1c**). Importantly, no equivalent density was observed in the sample where calcium was not included and divalent cations were chelated by EDTA (**Fig. 1c**, ‘No divalent cations’, **Extended Data Fig. 2**a**-c**), indicating that the observed cryo-EM densities represent Ca^2+^ ions.

**Figure 1.**
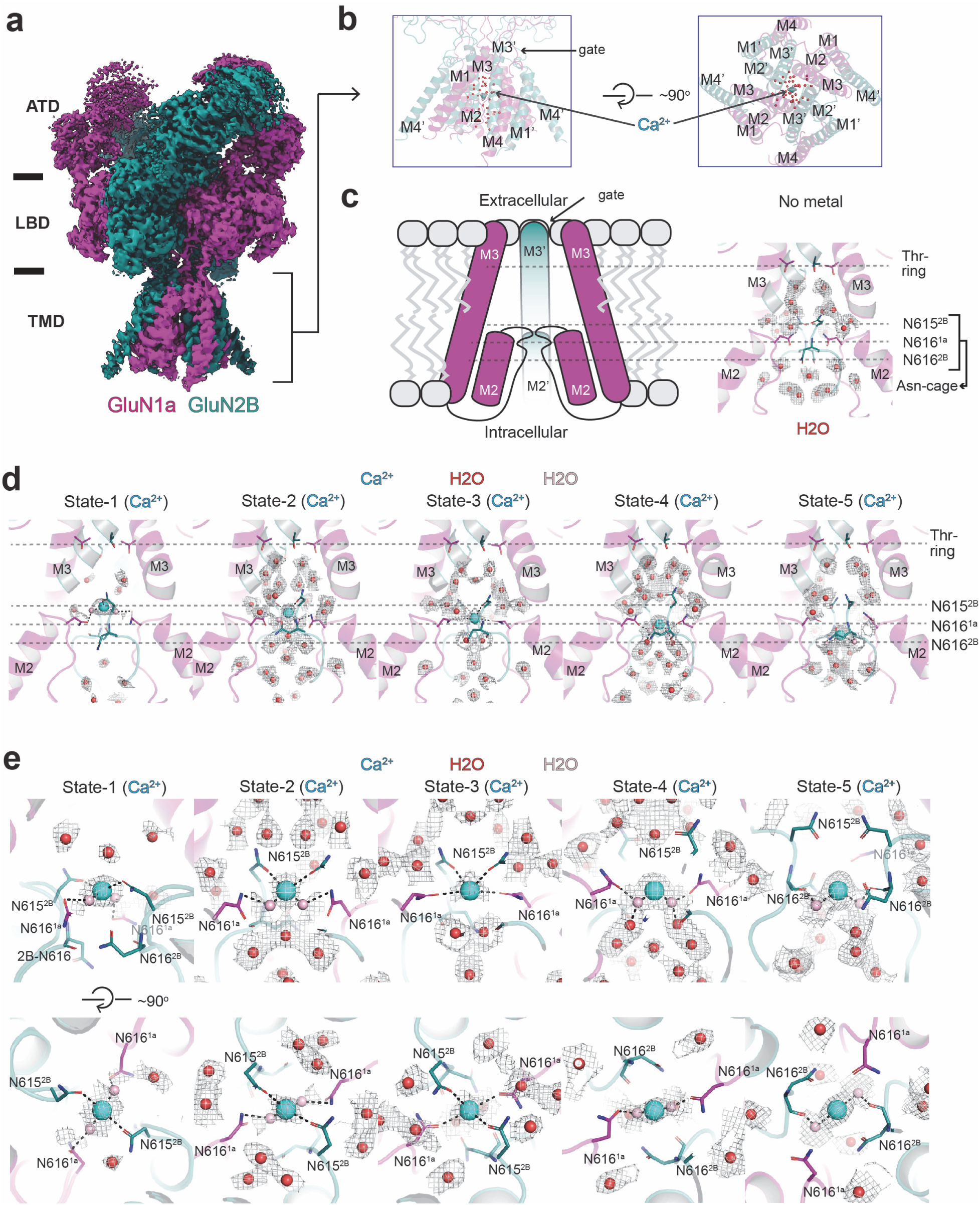
Structure of GluN1a-2B NMDAR in complex with agonists and calcium ions. **a-b,** Cryo-EM density (a) and the TMD model (b) of the GluN1a-2B NMDAR bound to agonists and Ca^2+^. One of the five Ca^2+^ (cyan sphere) positions and waters (red spheres) in the pore are shown. **c-d,** Schematic figure of the pore region (c) and a zoom-in view of the agonists-bound GluN1a-2B NMDAR with no divalent cation (EDTA-condition) showing cryo-EM density (gray mesh) for waters. The Thr-ring is a part of the channel gate, whereas Asn-cage constitutes the cationic selectivity filter. **d,** The views of the channel pore with Ca^2+^ at five distinct positions (State-1-5) from the top to the bottom of the Asn-cage. **e,** Close-up views of the Ca^2+^ sites viewed from the side and top of the Asn-cage. Pink spheres represent the putative water molecules directly coordinating Ca^2+^, modeled based on continuous density within 2.5 Å of Ca^2+^. Dotted lines represent polar interactions.

The Ca^2+^ ions occupy five positions within the Asn-cage, suggesting the permeation pathway (**Fig. 1d-e**). Here, we arbitrarily name them State-1 through 5, from the extracellular to the intracellular side. In State-1, 2 and 3, Ca^2+^ ions are located around the upper entry site of the Asn-cage, constituted by the two GluN2B Asn615 residues, forming the narrowest constriction within the NMDAR channel. The distances between Ca^2+^ and GluN2B Asn615 residues are sufficiently short to allow direct interactions, with Ca^2+^ being partially dehydrated (**Fig. 1d-e, State-1-3, dotted lines**). We speculate, based on continuous density with Ca^2+^ and interacting residues, that the hydration shell waters (pink spheres) make secondary contact with GluN1a Asn616 as well (**Fig. 1d-e, State-1-2**), whereas in State-3, Ca^2+^ is positioned to form the direct interactions with both GluN2B Asn615 and GluN1a Asn616 (**Fig. 1d-e, State-3**). In State-4 and 5, the interactions are through secondary interactions with the hydration shells and GluN1a Asn616 (State-4) or GluN2B Asn 616 main chain carbonyl (State-5) (**Fig. 1d-e, State-4-5**). Above and beneath the Asn-cage motif, there is sufficient space for Ca^2+^ to diffuse; therefore, no defined Ca^2+^ density was observed. Based on the structural observations of Ca^2+^ in various positions, we speculate that the partial dehydration of the Ca^2+^, mainly by GluN2B Asn615, is required for Ca^2+^ to pass through the Asn-cage (**Fig. 1**).

While we focused on analyses of the TMD region, the most predominant overall protein conformation here is the non-active state where glycine and glutamate are bound to GluN1 LBD and GluN2B LBD, respectively, while the channel gate at the entrance of the TMD channel is closed (Chou et al., 2020; Tajima et al., 2016). Our recent studies have shown that the channel pores surrounding the Asn-cage, located below (toward the cytoplasmic domain), remain largely unchanged between open and closed channels, remaining occupied with ions (Chou et al., 2024; Kang et al., 2025). Therefore, while our structure shows a closed gate at the extracellular entrance, the cation permeation pathway remains intact, justifying the calcium permeation mechanism through the Asn-cage.

### Two magnesium binding sites within the NMDAR channel pore

To understand the mechanism of channel blockade by Mg^2+^, we obtained the cryo-EM structure of the GluN1a-2B NMDAR in the presence of agonists (glycine and glutamate) and MgCl_2_ (**Fig. 2a- b**). We captured the structure in the agonist-bound form, allowing Mg^2+^ entry from the extracellular side, which represents the physiological pathway. The extracellular Mg^2+^ entry depends on the opening of the channel gate located at the juxtamembrane region (Chou et al., 2024; Kang et al., 2025) (**Fig. 2b**). Here, we implemented a similar strategy in single-particle analysis, including extensive 3D classification followed by a local refinement of the TMD to attain resolution at the TMD ranging from 2.7 to 3.0 Å to monitor Mg^2+^ (**Extended Data Fig. 2**d**-g**). Unlike the Ca^2+^-bound structures, there are strictly two distinct binding sites, one above the Asn-cage (upper) and the other at the intracellular side of the Asn-cage (Lower) (**Fig. 2c-d**). These densities are absent in the sample without added divalent metals (**Fig. 1c**) and are distinct from the Ca^2+^- bound structures. The upper site Mg^2+^ does not directly contact any residues as the closest residues, GluN2B Asn615 (side chain oxygens), are 4.1 Å away. Instead, the hydrated Mg^2+^ is in contact with the surrounding water molecule network, forming contact with GluN2B Asn615 (**Fig. 2c**). The Mg^2+^ hydration shell further interacts with surrounding waters that form a hydrogen bond network with the GluN1a Asn616 residues (**Fig. 2c-d, Upper**). Therefore, the binding of the extracellular Mg^2+^ involves a water-mediated network with the N-site, GluN1a Asn616, and GluN2B Asn615 (**Fig. 2c-d**). It is important to note that, unlike Ca^2+^, Mg^2+^ does not appear to traverse the Asn-cage, as no positions analogous to the multiple Ca^2+^ sites are observed. It has been well-documented that the dehydration of Mg^2+^ requires substantially more energy than that of Ca^2+^ due to the smaller ionic radius of Mg^2+^, which leads to a stronger attraction to the oxygen atoms of water and higher hydration energy (Ikeda et al., 2007). Since passing through the narrow Asn-cage would require dehydration, we suggest a higher dehydration energy requirement for Mg^2+^ disfavors the permeation and, instead, makes Mg^2+^ a blocker right above the two GluN2B Asn615 residues. Consistent with our structural observation as well as previous reports (Burnashev et al., 1992b; Wollmuth et al., 1998a), the GluN1a Asn616Gln and GluN2B Asn615Gln mutations robustly affect the voltage-sensitive Mg^2+^ block (**Fig. 3d, Extended Data Fig. 3)**.

**Figure 2.**
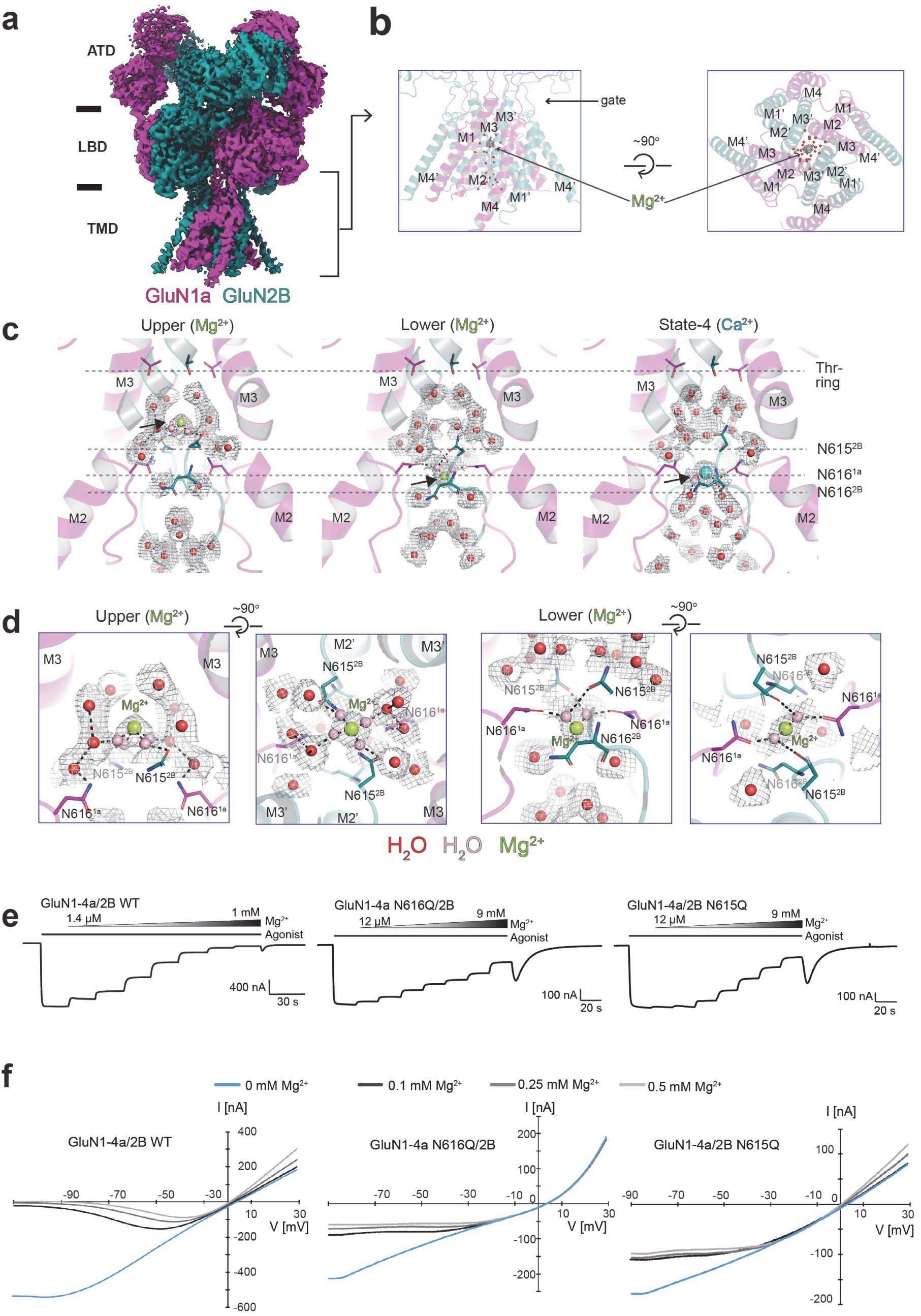
Structure of GluN1a-2B NMDAR in complex with agonists and magnesium ions. **a-b,** Cryo-EM density and the model showing the upper Mg^2+^ (limon sphere) and waters (red spheres) in the pore. **c,** The view of the channel pore in complex with two Mg^2+^ sites, Upper and Lower, in comparison with Ca^2+^ at the State-4 position (arrows). **d,** Zoom-in view of the Upper and Lower Mg^2+^ binding sites viewed from the side and top of the Asn-cage. The pink spheres represent the plausible directly coordinating waters with Mg^2+^, modeled based on continuous density with Mg^2+^. Dotted lines represent polar interactions. **e,** Concentration-responses of Mg^2+^ on wild-type and Asn-cage mutants at -60 mV holding potential. **f,** I-V curves at different Mg^2+^ concentrations. The recordings were done using TEVC on cRNA-injected oocytes. All currents were induced by the application of glycine and glutamate at 100 µM.

**Figure 3.**
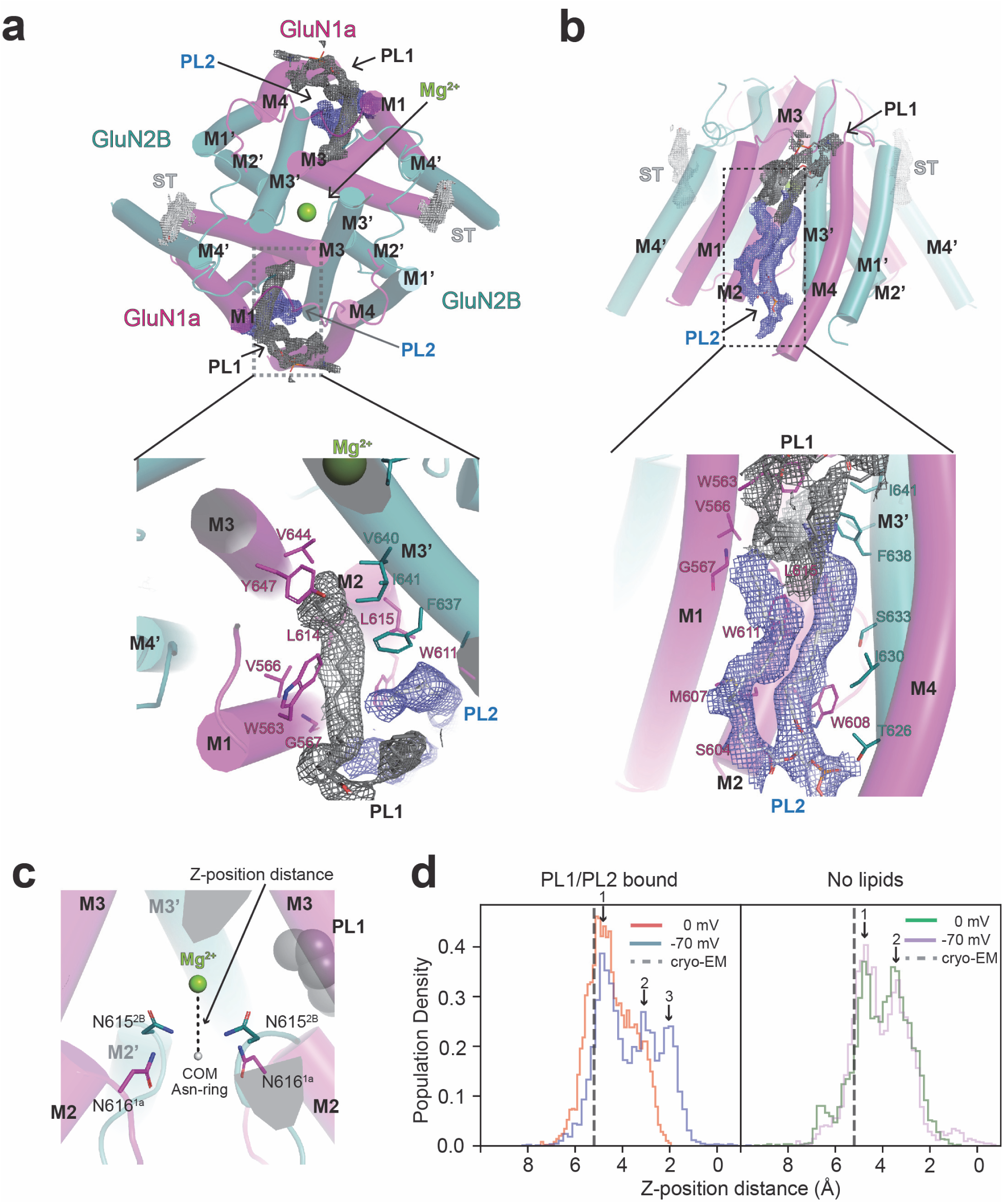
Lipid binding sites in GluN1a-2B NMDAR. **a-b,** Cryo-EM captures the tightly bound lipids in the TMD. Cryo-EM density for sterol-like density (ST, white mesh) around GluN2B M4’ and phospholipids 1 and 2 (PL1, dark gray mesh, PL2, slate mesh) viewed from the extracellular side (a) and side of the membrane (b). Note that one of the acyl chains of PL1 wedges into the back of the upper Mg^2+^ binding site (limon). **c-d,** MD simulations estimating the population density of Z-distance distribution between Mg^2+^ and the center of mass of GluN1 Asn616 and GluN2B Asn615 Cα atoms (**c**) at 0 mV and -70 mV in the presence (**d**, left) and absence (**d**, right) of PL1 and PL2. One of the stable positions identified by MD simulations, peak 1, corresponded closely to the cryo-EM-derived position (dashed line). At −70 mV, additional peaks (peaks 2 and 3) emerged at shorter Z-distances relative to 0 mV, suggesting voltage-dependent alterations in Mg²⁺ dynamics in the presence of PL1 and PL2 (**d**, left panel). By contrast, no such voltage-dependent shift was observed in the absence of these lipids, with population density distributions appearing similar between 0 mV and −70 mV (**d**, right panel). A bin width of 50 was used for each histogram.

The Mg^2+^ at the lower site indirectly interacts with GluN1 Asn616 and GluN2B Asn616 via a water network (**Fig. 2c-d, Lower**). This Mg^2+^ binding position is similar to State-4 in the Ca^2+^ structure (**Fig. 2c, State-4**), although the binding mode differs from each other (**Fig. 1c-d, State-4, Fig. 2c-d, Lower**). Given that the hydrated Mg^2+^ does not pass through the Asn-cage from the extracellular entrance, we speculate that the lower site represents the intracellular Mg^2+^ block site, which is accessed from the intracellular side (Johnson and Ascher, 1990). As our structure shows, the region outside the Asn-cage toward the intracellular side is wide open; therefore, Mg^2+^ has free access up to the Asn-cage in the context of our purified proteins. The lower Mg^2+^ site is also similar to the one observed in the antagonist-bound NMDAR (Huang et al.) (PDB code: 9YIP). Consistently, the upper Mg^2+^ site was not observed in the antagonist-bound NMDAR where the channel gate is strictly closed; therefore, there is no access to Mg^2+^ or any ions from the extracellular side. With respect to the lower Mg^2+^ site, the major difference in our structure is that the hydrated Mg^2+^ is making water-mediated network with GluN1a Asn616 and GluN2B Asn615 rather than the direct contact, which involves dehydration of Mg^2+^ (**Fig. 3d**). Nevertheless, our observation of the distinct sites for extracellular and intracellular Mg^2+^ sites is consistent with the previous suggestions based on additive effects of the two in electrophysiological experiments (Johnson and Ascher, 1990). While the extracellular Mg^2+^ channel blockade is more relevant in neuroplasticity, the intracellular Mg^2+^ block may modulate the excitability of neurons.

### Lipid network surrounds the NMDAR pore and affects voltage-dependent Mg^2+^ sensitivity

Our workflow in single-particle cryo-EM unambiguously captured the binding of phospholipids. They were carried over from the expression host cells since no lipids were added during the protein purification. In structures in all three conditions, no added divalent cation, Ca^2+^, and Mg^2+^, two lipids, PL1 and PL2, networked with one another, are observed at each GluN1a-GluN2B interface in the tetramer symmetrically (**Fig. 3a-b**). These lipids are located near the ‘back’ of the Asn-cage where the Mg^2+^ block occurs (**Fig. 3a-b**). One acyl chain from PL1 is placed almost parallel to the membrane plane and is surrounded by GluN1a M1, M2, and M3 and GluN2B M3’, and the other acyl chain is placed next to GluN1a M4 (**Fig. 3a**). The two acyl chains of PL2 are placed orthogonal to the membrane plane and are surrounded by GluN1a M1 and M2 and GluN2B M3’ (**Fig. 3b**). One of the two PL1 acyl chains wedges into a pocket and interacts with GluN1a Leu615, Trp611, Met607, Ser604, and GluN2B Phe637, and Ile641 (**Fig. 3a**). Residues such as GluN1a Met607, Ser604, Trp611, and GluN2B Ile630 and Ser633 interact with PL2 (**Fig. 3b**). PL1 and PL2 are tightly associated with each other and integrated into the protein structure as a single entity (**Fig. 3a-b**). Due to their proximity to the ‘back’ of the Mg^2+^ binding site, we hypothesized that these tightly bound lipids may regulate Mg^2+^ block at the Asn-cage region.

To explore this possibility, we analyzed the binding stability of the upper Mg^2+^ responsible for the voltage-dependent Mg^2+^ binding by calculating the distance between Mg^2+^ and the center of geometry of the c-α atoms of the Asn-ring (GluN1a Asn616 and GluN2B Asn615) in the presence and absence of lipids (PL1 and PL2) and at resting (-70 mV) and depolarizing (0 mV) membrane voltages by MD simulations (**Fig. 3c-d**). While the branching acyl chains of the phospholipids are visible, there was insufficient cryo-EM density to identify the headgroup definitively. Here, phosphatidylcholine was arbitrarily modeled as lipids. Our simulations showed that Mg^2+^ can move on the Z-axis but not pass the entrance of the Asn-cage, consistent with the notion that Mg^2+^ blocks the channel (**Fig. 3d**). In the presence of lipids, Mg^2+^ can more readily move closer to the Asn-cage at -70 mV (blue) than 0 mV (orange), indicative of preference of Mg^2+^ binding at resting potential (**Fig. 3d**, left panel, arrows). In the absence of lipids, the distribution of Mg^2+^ positions at 0 and -70 mV is similar, suggesting little or no voltage sensitivity (**Fig. 3d, right**). Thus, our MD simulations support the involvement of bound lipids in the voltage-sensitive Mg^2+^ block.

### Mutations on lipid-binding residues alter Mg^2+^ sensitivity

To further assess the involvement of PL1 and PL2 in voltage-dependent extracellular Mg^2+^ channel block, we incorporated site-directed mutations into the lipid binding sites (**Fig. 4a**, mutated residues are highlighted by ovals) and evaluated their effects on channel blockade using two-electrode voltage clamp (TEVC) electrophysiology (**Fig. 4b-e**). These point mutants were designed to perturb the binding of PL1, PL2, or both. Concentration responses at three different membrane voltages, -60, -40, and -20 mV, were measured and compared to the wild-type GluN1a-2B NMDAR (**Fig. 4b-e, Extended Data Table 2, Extended Data Fig. 4**). The effects of all GluN1a mutants were lowering of Mg^2+^ potency and could be grouped into three categories, significant changes at all voltages (Gly567Trp, Met607Trp; cyan in **Fig. 4a, c**), significant effect at -60 and -40 mV but not at -20 mV (Val566Trp and Leu615Gln, dark orange in **Fig. 4a, c**), and no effect at -60 and -40 mV and significant effect at -20 mV (Ser604Trp, gold in **Fig. 4a,c**). Furthermore, Leu615Trp did not exhibit any significant effect at any voltage (**Fig. 4b-c**). For GluN2B, Ser633Leu at all voltages had a considerable lowering of Mg^2+^ potency (cyan in **Fig. 4a,d**), Ile630Trp at -20 mV, but not at -60 mV and -40 mV, showed a significant effect (gold in **Fig. 4a, d**) whereas Thr626Trp and Phe637Trp had no significant effects (**Fig. 4d-e**). The mutations that target PL2 or both PL1 and 2 tend to affect Mg^2+^ potency at all voltages (1a-Gly567Trp, 1a- Met607Trp, and 2B Ser633Leu, cyan in **Fig. 4a, c, d**). The mutations near the head group of PL2 (1a-Ser604Trp and 2B-Ile630Trp, gold in Fig. 4a, c, d) tend to affect Mg^2+^ only at more depolarizing potentials, -20 mV. The mutations that only target PL1 (dark orange in **Fig. 4a, c, d**) tend to affect Mg^2+^ at -60 and -40 mV but not at -20 mV. The GluN2B Ser633Leu mutation was previously reported to alter Mg^2+^ potency (Siegler Retchless et al., 2012). It is essential to note that GluN2B Ser633 is located at the PL2 binding site, and we anticipate that the GluN2B Ser633Leu mutant interacts with PL2 differently. Overall, the cryo-EM, MD simulations, and electrophysiology data presented above indicate that lipids play a crucial role in controlling Mg^2+^ block.

**Figure 4.**
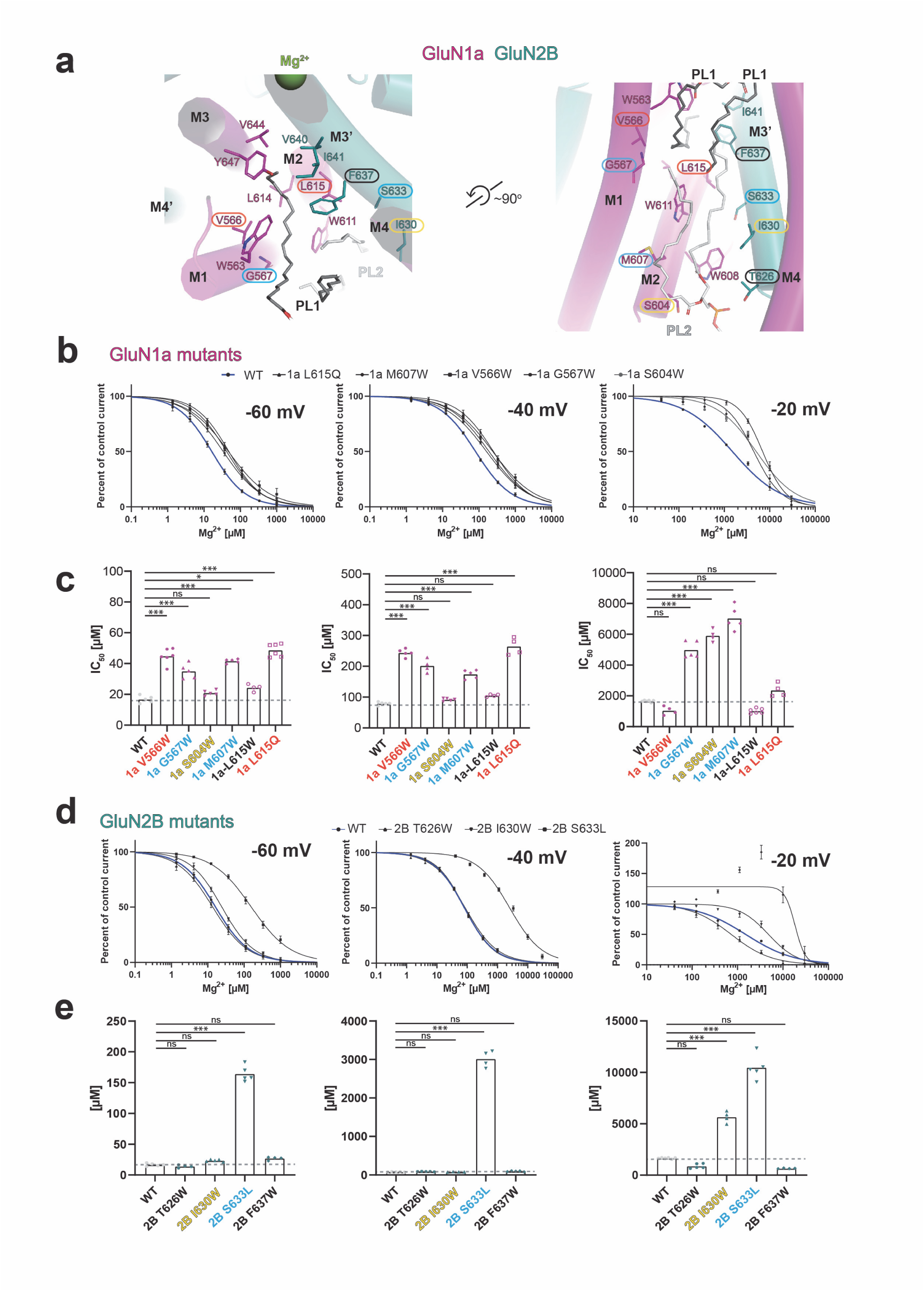
Effects of lipids on voltage-dependent Mg^2+^ block in GluN1a-2B NMDAR. **a,** Close-up views of PL1 (dark gray sticks) and PL2 (white sticks) binding sites from the top (left) and side (right) of the membrane. Residues enclosed by cyan, orange, and yellow ovals have altered Mg^2+^ sensitivity when mutated at all tested voltages, at -60 and -40 mV, and only at -20 mV, respectively. The residues outlined in black show no mutational effect at any voltage. **b-d,** Mg^2+^ concentration-response curves (**b, d**) and corresponding IC_50_ values (**c, e**) derived from the TEVC recording at -60, -40, -20 mV for the GluN1a (**b-c**) and GluN2B mutants (**d-e**). The color codes of the mutants are the same as in panel a. The isotype of GluN1a used in these TEVC experiments is GluN1-4a. The statistical analysis was performed by One-Way ANOVA (*** p < 0.001, ** 0.001 < p < 0.01, * 0.01 < p < 0.05, n.s. not significant.

## Discussion

The Ca^2+^ permeation and voltage-dependent channel blockade by extracellular Mg^2+^ are two fundamental molecular properties of NMDARs that are crucially involved in neural plasticity. Our cryo-EM structures captured Ca^2+^ at five positions throughout the cation selectivity filter (Asn- cage) and Mg^2+^ at two distinct positions above and in the lower part of the Asn-cage. Based on our cryo-EM structures, we propose that Ca^2+^ permeability is facilitated by the ability of the NMDAR Asn-cage to directly coordinate with and partially dehydrate Ca^2+^ (**Fig. 5a**). In contrast, the NMDAR Asn-cage is unable to dehydrate Mg^2+^, a process that incurs a substantially higher

**Figure 5.**
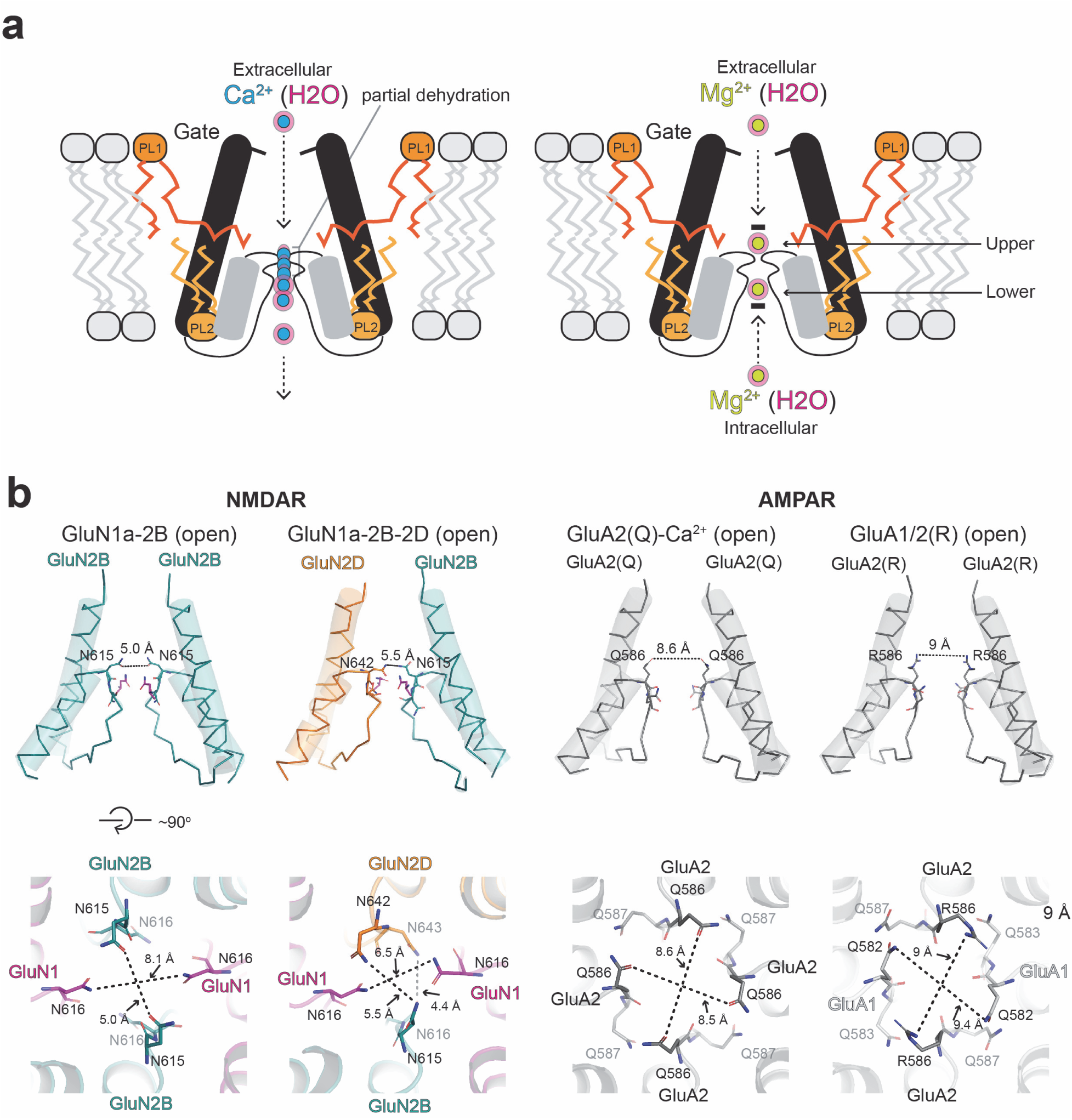
Molecular mechanism of Ca^2+^ permeability and Mg^2+^ block in NMDAR and comparison with AMPARs. **a,** Cation permeation through the narrow Asn-cage requires partial dehydration. Hydrated Ca^2+^ (concentric cyan and light pink) can undergo partial dehydration as it permeates through the Asn-cage. In contrast, Mg^2+^ requires substantially higher energy for dehydration, making its permeation through Asn- cage energetically unfavorable. Instead, hydrated Mg^2+^ (concentric limon and light pink) binds at upper and lower sites via a structured water network, corresponding to the extracellular and intracellular Mg^2+^ block sites, respectively. Note that extracellular Mg^2+^ can access the upper site only when the channel gate is open in response to agonists. The voltage sensitivity of the extracellular Mg^2+^ block may be partly regulated by the tightly bound lipids, PL1 and PL2 (dark and light orange). **b,** Side views of open channel structures for GluN1a-2B (PDB code: 9ARE), GluN1a-2B-2D (PDB code: 9D38), GluA2(Q)/TARP gamma-2 (PDB code: 8FP9), and GluA1/2 (R)/TARP gamma-8 (PDB code: 7OCF). TARP gamma-2 and -8 are omitted for visual comparison with NMDARs. Note that the pore dimensions of the QRN sites are substantially wider in AMPARs than in NMDARs, suggesting a possibility that Ca^2+^ permeation may occur without dehydration in AMPARs.

energetic cost compared to Ca^2+^. Consequently, Mg^2+^ cannot traverse the Asn-cage and instead forms a water-mediated network that enables it to act as a channel blocker (**Fig. 5a**). It is worth noting that this was presciently proposed by Ascher and Nowak in 1988, well before the structures of any channel were available (Ascher and Nowak, 1988). In the current study, we identified two Mg^2+^ binding sites, upper and lower, which likely correspond to the extracellular and intracellular Mg^2+^ blockade sites, respectively. The presence of agonists in our cryo-EM samples, which promotes channel opening (Chou et al., 2024; Kang et al., 2025), allows Mg^2+^ ions to access the Asn-cage from both extracellular and intracellular directions. In contrast, a recently published Mg^2+^ bound structure in the presence of an antagonist (Huang et al., 2025) captured a single Mg^2+^ at a site similar to the ‘Lower’ site but not the ‘Upper site.’ This is consistent since the occupancy of the upper site requires the opening of the channel gate, which, in the antagonist-bound structure, is closed (Chou et al., 2020) and, therefore, shuts down the entry of extracellular Mg^2+^. Nevertheless, the distinct existence of the extracellular and intracellular Mg^2+^ sites aligns with previous electrophysiological findings where the blocking effects of Mg^2+^ from either side were additive (Johnson and Ascher, 1990; Wollmuth et al., 1998b). The physiological relevance of the intracellular Mg^2+^ block remains unclear.

The architecture of the QRN site in NMDAR allows Ca^2+^ to become partly dehydrated, conferring a Ca^2+^ selectivity. In addition, there are extracellular factors, including the DRPEER motif (Watanabe et al., 2002), shown to regulate Ca^2+^ permeability and selectivity. It would be meaningful to explore whether the mechanism of Ca^2+^ permeability in NMDAR and the CP- AMPAR is similar. A recent study on the open-state CP-GluA2 AMPAR (Q)/TARP gamma-2 revealed that Ca^2+^ is captured around the channel gate (G-site) but not in the QNR site (Nakagawa et al., 2024), potentially indicating the absence of metastable Ca^2+^ binding in its selectivity filter. The channel pores of CP-GluA2 AMPAR/TARP gamma-2 (Twomey et al., 2017) and GluA1/2 AMPAR/TARP gamma-8 (Zhang et al., 2021) in the open state are substantially larger in diameter than the open state GluN1a-2B (Chou et al., 2024) and GluN1a-2B-2D NMDAR (Kang et al., 2025) (**Fig. 5b, Extended Data Fig. 5**). Consistent with this hypothesis, Mg^2+^, with a smaller hydration shell dimension than Ca^2+^, has been reported to permeate through CP-GluA2 (Burnashev et al., 1992a). In contrast, the GluN1a-2B NMDAR QNR site is more constricted than AMPAR to disallow the passing of Ca^2+^ without dehydration (**Fig. 5b**). Therefore, our structural observations propose the possibility that the mechanism of calcium selectivity differs between CP-AMPAR and NMDAR, which could explain higher Ca^2+^ selectivity over Na^+^ for NMDARs (Wollmuth and Sakmann, 1998). The dimension of the QNR site is even smaller in the GluN1a-2B-2D NMDAR (**Fig. 5b**), consistent with the previous report that the inclusion of GluN2D lowers Ca^2+^ permeability (Kuner and Schoepfer, 1996). Finally, it is essential to note that the Asn-cage residues in GluN2A and GluN2B are susceptible to *de novo* mutations in humans, which alter voltage-dependent Mg^2+^ block and are consequently associated with neurological diseases, including autism spectrum disorder, epilepsy, intellectual disability, and developmental delay (Li et al., 2019), indicating the crucial role of the Asn-cage in neuronal functions in brain development.

## Acknowledgment

We thank N. Simorowski for technical support. D. Thomas and M. Wang are thanked for managing the cryo-EM facility and computing facility at Cold Spring Harbor Laboratory, respectively. We thank T. Nakagawa for their critical comments on this work. This work was funded by the NIH (NS111745 and MH085926 to H.F), Austin’s purpose, Robertson funds at CSHL, Doug Fox Alzheimer’s fund, Heartfelt Wing Alzheimer’s fund, and the Gertrude and Louis Feil Family Trust (all to H.F). The computational work was performed with assistance from an NIH grant (S10OD028632-01)

## Contributions

R.S., T-H.C., and H.F. conceived the project. R.S. and T-H.C. obtained all cryo-EM structures. R.S. conducted electrophysiology experiments. M.E. conducted MD simulations. R.S. and H.F. wrote the manuscript with input from all authors.

## Methods

### Expression and purification of GluN1a-2B NMDAR

The rat GluN1a-2B NMDAR was expressed using the EarlyBac system (Furukawa et al., 2021) and purified using the previously established method (Regan et al., 2018). Sf9 insect cells at 4.0 × 10^6^ cells/mL were infected with the recombinant EarlyBac baculovirus harboring both GluN1a and CTD truncated GluN2B (residue 27–852 N-terminally tagged with a dual strep-tag after the *Xenopus* GluN1 signal peptide and with Cys849Ser. To improve the expression level, 6 of 11 glycosylation sites in GluN1a were mutated as follows: Asn61Gln, Asn239Asp, Asn350Gln, Asn471Gln, Asn491Gln, and Asn771Gln. Finally, the ER retention signal (Arg/Arg/Lys) at the GluN1a construct was altered by the mutations Arg844Gln, Arg845Gly, and Lys846Ala. Cells were harvested after 48 hours of infection and resuspended in 20 mM HEPES-Na pH 7.5, 150 mM NaCl, 1 mM glycine, 1 mM Na-glutamate, and 1 mM phenylmethylsulfonyl fluoride (PMSF). Lysis was performed with the aid of Emulsiflex C3 (Avestin). The membrane fraction was obtained through centrifugation at 40,000 rpm at 4 °C for 30 min. The membrane-containing pellet was resuspended to a concentration of 100 mg/mL in buffer containing 20 mM HEPES-Na pH 7.5, 150 mM NaCl, 1 mM glycine, 1 mM Na-glutamate, and 0.5 % lauryl maltose neopentyl glycol (LMNG). The desired protein was solubilized for 2 hours at 4 °C. Insoluble material was removed through centrifugation at 40.000 rpm for 30 minutes at 4 °C. Solubilized Strep-tagged GluN1a- 2B NMDAR was purified from the supernatant using Strep-tactin Sepharose, followed by size-exclusion chromatography (SEC, Superose 6 Increase column from GE Healthcare) in buffer containing 20 mM HEPES-Na pH 7.5, 150 mM NaCl, 0.002% LMNG, 1 mM glycine, and 1 mM Na- glutamate. NMDAR containing fractions were concentrated to a protein content of 1-4 mg/mL. The cation-free NMDAR sample was obtained by adding 1 mM EDTA to each buffer during purification. For the cation-bound structures, the ions (10 mM Ca^2+^ and 100 mM Mg^2+^) were added to the purified protein, and the mixture was incubated on ice for 30 minutes.

### Single-particle cryo-EM analysis on cation-bound GluN1a-2B NMDARs

The cation-free, and Ca-bound NMDAR samples were blotted on Quantifoil R 1.2/1.3 + 2 nm C grids (Quantifoil), and Mg-bound NMDAR samples were blotted on UltrAufoil holey gold film grids (Quantifoil) using Leica GP2 at 18 °C and at 85% humidity (blot time = 2.0 - 3.0 seconds) and vitrified in liquid ethane. Micrographs were acquired by Titan Krios (FEI) at Cold Spring Harbor Laboratory, operating at 300 keV, and the GATAN K3 Summit direct electron detector coupled with the GIF quantum energy filter (Gatan) at ×105,000 magnification (0.827 – 0.856 Å/pixel), defocus range of −2.4 to −0.6 µm, 30 or 40 frames, and 2- to 2.8-second exposure totaling a dose ranging 55.6–71.7 e^−^/Å^2^. All data sets were processed using Cryosparc v4.2.1 or v4.4.1 (Punjani and Fleet, 2021). Collected movies were motion corrected and CTF-estimated. For particle picking, templates were made from WARP-processed micrographs. Particles were picked using a particle size of 180 Å. Picked particles were extracted and cropped in a 4x box size to enhance the processing speed. Extracted particles were passed through multiple rounds of 2D classification, ab-initio, and heterogeneous refinement to reduce the amount of junk particles. Final particle sets were re-extracted to 400 - 440 box size and refined using non-uniform refinement. 3D classification was used to distinguish between different states. To enhance the overall resolution and local motion correction, the particles were further refined using non-uniform refinement and local refinement (or Reference-based motion correction). Model building was initially done by docking GluN1a-2B (PDB: 7SAA) to the cryo-EM density map using ChimeraX v1.4 (Meng et al., 2023). Further processing was done using Phenix v1.19.2-4158 (Liebschner et al., 2019) and manually refined using COOT v.0.9.8.1 (Emsley and Cowtan, 2004). A summary of data collection and refinement statistics is shown in **Extended Data Table 1**.

### Equilibrium MD simulations

Equilibrium MD simulations were performed using OpenMM v7.5.1 (Eastman et al., 2017). Temperature was set to 298 K using the Langevin Middle integrator with friction coefficient of 1 ps. Pressure was maintained at 1 bar with the Monte Carlo anisotropic barostat. Electrostatics were computed with Particle Mesh Ewald, with a non-bonded cutoff of 1 nm. Hydrogen masses were repartitioned to 1.5 times their typical molecular weight, allowing for a larger integration step size of 4 fs to be implemented. The protein model consisted of both LBD and TMD and without Cα atom position restraints. For each system, a total of 10 x 200 ns independent simulations were performed.

### Electrophysiology

To express di-heteromeric NMDARs in *Xenopus laevis* oocytes for two-electrode voltage clamp (TEVC) experiments, we used rGluN1, rGluN2B DNA constructs, as previously described (Karakas et al., 2009). cRNAs were transcribed using mMessage mMachine t7 transcription kit (Invitrogen) and subsequently injected into defolliculated *Xenopus laevis* oocytes (Ecocyte Bioscience) with a total amount of up to 25 ng. Injected oocytes were further incubated in 50% L-15 medium supplemented with 15 mM HEPES pH = 7.5, 100x pen-strep solution (Thermofisher), and 3 % FBS for 1-2 days at 18 degrees. TEVC (Axoclamp-2B) recordings were performed using an extracellular solution containing 5 mM HEPES, 100 mM NaCl, 0.3 mM BaCl2, and 10 mM Tricine at a final pH of 7.4 (adjusted with KOH). The current was measured using an agarose-tipped microelectrode (0.4–0.9 MU) at the holding potential of -90 to -20 mV. Maximal response currents were evoked by 100 µM of glycine and 100 µM of L-glutamate. Data were acquired by the PatchMaster program (HEKA) and analyzed using Origin 8 (OriginLab Corp).

### Quantification and statistical analysis

Statistical analysis of TEVC and whole-cell patch clamp was done using Graphpad Prism and can be found in the corresponding figures and figure legends. Error bars in the fitted curve represent mean ± SD. N represents the number of independent recordings done on separate cells.

## Data Availability

Cryo-EM density maps have been deposited in the Electron Microscopy Data Bank (EMDB). The structural coordinates have been deposited in the RCSB Protein Data Bank (PDB).

## References

Ascher, P., and Nowak, L. (1988). The role of divalent cations in the N-methyl-D-aspartate responses of mouse central neurones in culture. J Physiol 399, 247-266.

Bliss, T.V.P., and Collingridge, G.L. (1993). A synaptic model of memory: long-term potentiation in the hippocampus. Nature 361, 31-39.

Burnashev, N., Monyer, H., Seeburg, P.H., and Sakmann, B. (1992a). Divalent ion permeability of AMPA receptor channels is dominated by the edited form of a single subunit. Neuron 8, 189–198.

Burnashev, N., Schoepfer, R., Monyer, H., Ruppersberg, J.P., Gunther, W., Seeburg, P.H., and Sakmann, B. (1992b). Control by asparagine residues of calcium permeability and magnesium blockade in the NMDA receptor. Science 257, 1415-1419.

Chou, T.H., Epstein, M., Fritzemeier, R.G., Akins, N.S., Paladugu, S., Ullman, E.Z., Liotta, D.C., Traynelis, S.F., and Furukawa, H. (2024). Molecular mechanism of ligand gating and opening of NMDA receptor. Nature 632, 209-217.

Chou, T.H., Tajima, N., Romero-Hernandez, A., and Furukawa, H. (2020). Structural Basis of Functional Transitions in Mammalian NMDA Receptors. Cell 182, 357-371 e313.

Eastman, P., Swails, J., Chodera, J.D., McGibbon, R.T., Zhao, Y., Beauchamp, K.A., Wang, L.-P., Simmonett, A.C., Harrigan, M.P., Stern, C.D., et al. (2017). OpenMM 7: Rapid development of high performance algorithms for molecular dynamics. PLOS Computational Biology 13, e1005659.

Emsley, P., and Cowtan, K. (2004). Coot: model-building tools for molecular graphics. Acta Crystallogr D Biol Crystallogr 60, 2126-2132.

Furukawa, H., Simorowski, N., and Michalski, K. (2021). Effective production of oligomeric membrane proteins by EarlyBac-insect cell system. Methods Enzymol 653, 3-19.

Hansen, K.B., Wollmuth, L.P., Bowie, D., Furukawa, H., Menniti, F.S., Sobolevsky, A.I., Swanson, G.T., Swanger, S.A., Greger, I.H., Nakagawa, T., et al. (2021). Structure, Function, and Pharmacology of Glutamate Receptor Ion Channels. Pharmacol Rev 73, 298-487.

Huang, X., Sun, X., Wang, Q., Zhang, J., Wen, H., Chen, W.-J., and Zhu, S. Structural insights into the diverse actions of magnesium on NMDA receptors. Neuron.

Huang, X., Sun, X., Wang, Q., Zhang, J., Wen, H., Chen, W.-J., and Zhu, S. (2025). Structural insights into the diverse actions of magnesium on NMDA receptors. Neuron 113, 1006–1018.e1004.

Ikeda, T., Boero, M., and Terakura, K. (2007). Hydration properties of magnesium and calcium ions from constrained first principles molecular dynamics. J Chem Phys 127, 074503.

Jain, A., Nakahata, Y., Pancani, T., Watabe, T., Rusina, P., South, K., Adachi, K., Yan, L., Simorowski, N., Furukawa, H., et al. (2024). Dendritic, delayed, stochastic CaMKII activation in behavioural time scale plasticity. Nature 635, 151-159.

Johnson, J.W., and Ascher, P. (1990). Voltage-dependent block by intracellular Mg2+ of N-methyl-D- aspartate-activated channels. Biophys J 57, 1085-1090.

Kang, H., Epstein, M., Banke, T.G., Perszyk, R., Simorowski, N., Paladugu, S., Liotta, D.C., Traynelis, S.F., and Furukawa, H. (2025). Structural basis for channel gating and blockade in tri-heteromeric GluN1-2B-2D NMDA receptor. Neuron.

Karakas, E., and Furukawa, H. (2014). Crystal structure of a heterotetrameric NMDA receptor ion channel. Science 344, 992-997.

Karakas, E., Simorowski, N., and Furukawa, H. (2009). Structure of the zinc-bound amino-terminal domain of the NMDA receptor NR2B subunit. The EMBO journal 28, 3910-3920.

Kuner, T., and Schoepfer, R. (1996). Multiple structural elements determine subunit specificity of Mg2+ block in NMDA receptor channels. J Neurosci 16, 3549-3558.

Kuner, T., Wollmuth, L.P., Karlin, A., Seeburg, P.H., and Sakmann, B. (1996). Structure of the NMDA receptor channel M2 segment inferred from the accessibility of substituted cysteines. Neuron 17, 343- 352.

Lee, C.H., Lu, W., Michel, J.C., Goehring, A., Du, J., Song, X., and Gouaux, E. (2014). NMDA receptor structures reveal subunit arrangement and pore architecture. Nature 511, 191-197.

Li, J., Zhang, J., Tang, W., Mizu, R.K., Kusumoto, H., XiangWei, W., Xu, Y., Chen, W., Amin, J.B., Hu, C., et al. (2019). De novo GRIN variants in NMDA receptor M2 channel pore-forming loop are associated with neurological diseases. Hum Mutat.

Liebschner, D., Afonine, P.V., Baker, M.L., Bunkoczi, G., Chen, V.B., Croll, T.I., Hintze, B., Hung, L.-W., Jain, S., McCoy, A.J., et al. (2019). Macromolecular structure determination using X-rays, neutrons and electrons: recent developments in Phenix. Acta Crystallographica Section D 75, 861-877.

Lisman, J., Malenka, R.C., Nicoll, R.A., and Malinow, R. (1997). Learning mechanisms: the case for CaM-KII. Science 276, 2001-2002.

MacDermott, A.B., Mayer, M.L., Westbrook, G.L., Smith, S.J., and Barker, J.L. (1986). NMDA-receptor activation increases cytoplasmic calcium concentration in cultured spinal cord neurones. Nature 321, 519- 522.

Malenka, R.C., and Bear, M.F. (2004). LTP and LTD: An Embarrassment of Riches. Neuron 44, 5-21.

Mayer, M.L., Westbrook, G.L., and Guthrie, P.B. (1984). Voltage-dependent block by Mg2+ of NMDA responses in spinal cord neurones. Nature 309, 261-263.

Meng, E.C., Goddard, T.D., Pettersen, E.F., Couch, G.S., Pearson, Z.J., Morris, J.H., and Ferrin, T.E. (2023). UCSF ChimeraX: Tools for structure building and analysis. Protein Science 32, e4792.

Miguez-Cabello, F., Wang, X.-t., Yan, Y., Brake, N., Alexander, R.P.D., Perozzo, A.M., Khadra, A., and Bowie, D. (2025). GluA2-containing AMPA receptors form a continuum of Ca2+-permeable channels. Nature.

Nakagawa, T., Wang, X.-t., Miguez-Cabello, F.J., and Bowie, D. (2024). The open gate of the AMPA receptor forms a Ca2+ binding site critical in regulating ion transport. Nature Structural & Molecular Biology 31, 688-700.

Nowak, L., Bregestovski, P., Ascher, P., Herbet, A., and Prochiantz, A. (1984). Magnesium gates glutamate- activated channels in mouse central neurones. Nature 307, 462-465.

Punjani, A., and Fleet, D.J. (2021). 3D variability analysis: Resolving continuous flexibility and discrete heterogeneity from single particle cryo-EM. Journal of Structural Biology 213, 107702.

Regan, M.C., Grant, T., McDaniel, M.J., Karakas, E., Zhang, J., Traynelis, S.F., Grigorieff, N., and Furukawa, H. (2018). Structural Mechanism of Functional Modulation by Gene Splicing in NMDA Receptors. Neuron 98, 521-529 e523.

Siegler Retchless, B., Gao, W., and Johnson, J.W. (2012). A single GluN2 subunit residue controls NMDA receptor channel properties via intersubunit interaction. Nat Neurosci 15, 406-413, S401-402.

Sommer, B., Köhler, M., Sprengel, R., and Seeburg, P.H. (1991). RNA editing in brain controls a determinant of ion flow in glutamate-gated channels. Cell 67, 11-19.

Tajima, N., Karakas, E., Grant, T., Simorowski, N., Diaz-Avalos, R., Grigorieff, N., and Furukawa, H. (2016). Activation of NMDA receptors and the mechanism of inhibition by ifenprodil. Nature 534, 63-68.

Twomey, E.C., Yelshanskaya, M.V., Grassucci, R.A., Frank, J., and Sobolevsky, A.I. (2017). Channel opening and gating mechanism in AMPA-subtype glutamate receptors. Nature 549, 60-65.

Watanabe, J., Beck, C., Kuner, T., Premkumar, L.S., and Wollmuth, L.P. (2002). DRPEER: a motif in the extracellular vestibule conferring high Ca2+ flux rates in NMDA receptor channels. J Neurosci 22, 10209- 10216.

Wollmuth, L.P., Kuner, T., and Sakmann, B. (1998a). Adjacent asparagines in the NR2-subunit of the NMDA receptor channel control the voltage-dependent block by extracellular Mg2+. J Physiol 506 ( *Pt 1*), 13-32.

Wollmuth, L.P., Kuner, T., and Sakmann, B. (1998b). Intracellular Mg2+ interacts with structural determinants of the narrow constriction contributed by the NR1-subunit in the NMDA receptor channel. J Physiol 506 ( *Pt 1*), 33-52.

Wollmuth, L.P., and Sakmann, B. (1998). Different mechanisms of Ca2+ transport in NMDA and Ca2+- permeable AMPA glutamate receptor channels. J Gen Physiol 112, 623-636.

Yasuda, R., Hayashi, Y., and Hell, J.W. (2022). CaMKII: a central molecular organizer of synaptic plasticity, learning and memory. Nature Reviews Neuroscience 23, 666-682.

Zhang, D., Watson, J.F., Matthews, P.M., Cais, O., and Greger, I.H. (2021). Gating and modulation of a hetero-octameric AMPA glutamate receptor. Nature 594, 454-458.

Zhou, C., and Tajima, N. (2023). Structural insights into NMDA receptor pharmacology. Biochem Soc Trans 51, 1713-1731.

