## Extended Data Figures and Tables for "On the mechanism of calcium permeability and magnesium block in NMDA receptors - a central molecular paradigm in neuroplasticity"

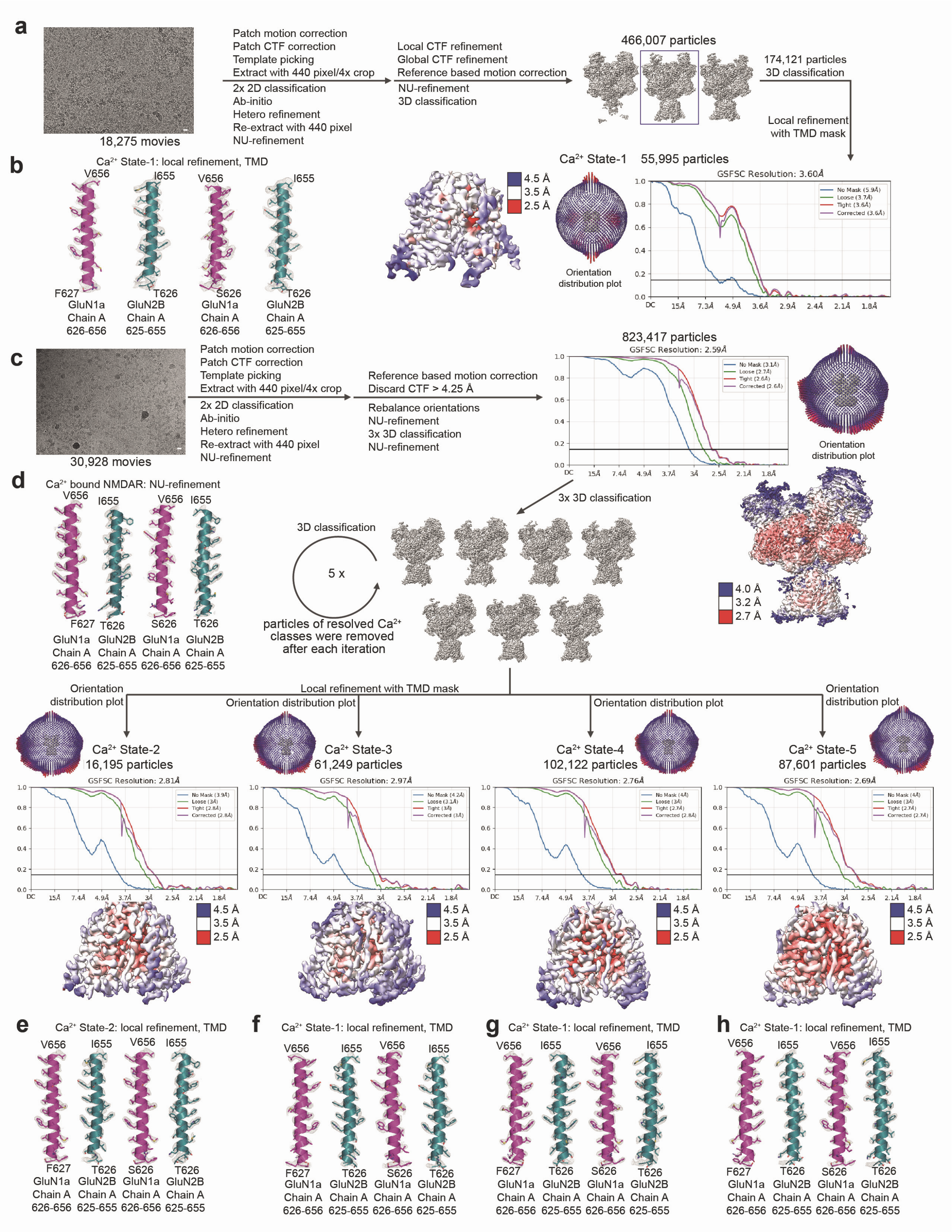
a
18,275 movies
Patch motion correction
Patch CTF correction
Template picking
Extract with 440 pixel/4x crop
2x 2D classification
Ab-initio
Hetero refinement
Re-extract with 440 pixel
NU-refinement
Local CTF refinement
Global CTF refinement
Reference based motion correction
NU-refinement
3D classification
466,007 particles
174,121 particles
3D classification
Local refinement
with TMD mask
b
Ca2+ State-1: local refinement, TMD
V656
I655
V656
I655
F627
T626
S626
T626
GluN1a Chain A 626-656
GluN2B Chain A 625-655
GluN1a Chain A 626-656
GluN2B Chain A 625-655
4.5 Å
3.5 Å
2.5 Å
Ca2+ State-1
55,995 particles
Orientation distribution plot
GSFSC Resolution: 3.60Å
c
30,928 movies
Patch motion correction
Patch CTF correction
Template picking
Extract with 440 pixel/4x crop
2x 2D classification
Ab-initio
Hetero refinement
Re-extract with 440 pixel
NU-refinement
Reference based motion correction
Discard CTF > 4.25 Å
Rebalance orientations
NU-refinement
3x 3D classification
NU-refinement
823,417 particles
GSFSC Resolution: 2.59Å
Orientation distribution plot
d
Ca2+ bound NMDAR: NU-refinement
V656
I655
V656
I655
F627
T626
S626
T626
GluN1a Chain A 626-656
GluN2B Chain A 625-655
GluN1a Chain A 626-656
GluN2B Chain A 625-655
3x 3D classification
3D classification
5 x
particles of resolved Ca2+ classes were removed after each iteration
4.0 Å
3.2 Å
2.7 Å
Local refinement with TMD mask
Orientation distribution plot
Ca2+ State-2
16,195 particles
GSFSC Resolution: 2.81Å
Orientation distribution plot
Ca2+ State-3
61,249 particles
GSFSC Resolution: 2.97Å
Orientation distribution plot
Ca2+ State-4
102,122 particles
GSFSC Resolution: 2.76Å
Orientation distribution plot
Ca2+ State-5
87,601 particles
GSFSC Resolution: 2.69Å
4.5 Å
3.5 Å
2.5 Å
e
Ca2+ State-2: local refinement, TMD
f
Ca2+ State-1: local refinement, TMD
g
Ca2+ State-1: local refinement, TMD
h
Ca2+ State-1: local refinement, TMD
V656
I655
V656
I655
F627
T626
S626
T626
GluN1a Chain A 626-656
GluN2B Chain A 625-655
GluN1a Chain A 626-656
GluN2B Chain A 625-655

**Extended Data Figure 1. Single-particle cryo-EM of the $Ca^{2+}$-bound GluN1a-2B NMDAR (Related to Figure 1).** In order to successfully obtain $Ca^{2+}$-bound GluN1a-2B NMDAR, iterative TMD-masked 3D classification without alignment was applied. Particles of well-resolved classes were separated after each iteration. The remaining particles underwent additional 3D classification until we stopped observing $Ca^{2+}$-bound classes. Each observed class was further evaluated through TMD-masked local refinement. **a,** Single-particle cryo-EM workflow of the $Ca^{2+}$ State-1 GluN1a-2B NMDAR. **b,** Map quality assessment of the $Ca^{2+}$ State-1 GluN1a-2B NMDAR at the M3/M3' region. **c,** Single-particle cryo-EM workflow of the $Ca^{2+}$ State-2-5 GluN1a-2B NMDAR. **d,** Map quality assessment of $Ca^{2+}$ bound GluN1a-2B NMDAR NU-refinement at the M3/M3' region. **e-h,** Map quality assessment of the $Ca^{2+}$ State-2-5 GluN1a-2B NMDAR at the M3/M3' region. The scale bars in the micrographs (panel a and c) represent 20 nm.

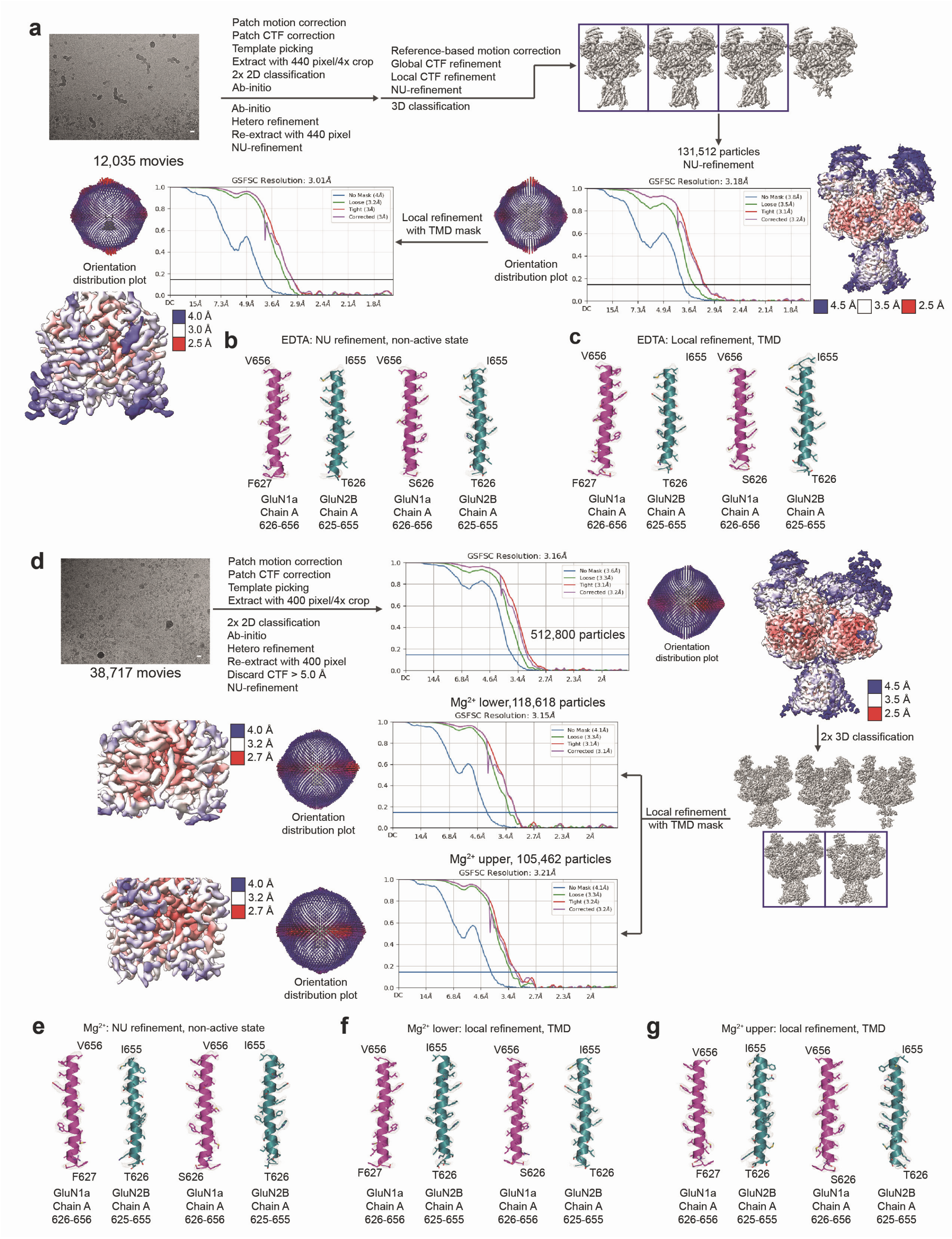
a
Patch motion correction
Patch CTF correction
Template picking
Extract with 440 pixel/4x crop
2x 2D classification
Ab-initio
Ab-initio
Hetero refinement
Re-extract with 440 pixel
NU-refinement
Reference-based motion correction
Global CTF refinement
Local CTF refinement
NU-refinement
3D classification
12,035 movies
131,512 particles
NU-refinement
GSFSC Resolution: 3.01Å
GSFSC Resolution: 3.18Å
Local refinement
with TMD mask
Orientation
distribution plot
Orientation
distribution plot
4.5 Å
3.5 Å
2.5 Å
4.0 Å
3.0 Å
2.5 Å
b
EDTA: NU refinement, non-active state
V656
I655
V656
I655
F627
T626
S626
T626
GluN1a
Chain A
626-656
GluN2B
Chain A
625-655
GluN1a
Chain A
626-656
GluN2B
Chain A
625-655
c
EDTA: Local refinement, TMD
V656
I655
V656
I655
F627
T626
S626
T626
GluN1a
Chain A
626-656
GluN2B
Chain A
625-655
GluN1a
Chain A
626-656
GluN2B
Chain A
625-655
d
Patch motion correction
Patch CTF correction
Template picking
Extract with 400 pixel/4x crop
2x 2D classification
Ab-initio
Hetero refinement
Re-extract with 400 pixel
Discard CTF > 5.0 Å
NU-refinement
38,717 movies
GSFSC Resolution: 3.16Å
512,800 particles
Orientation
distribution plot
4.5 Å
3.5 Å
2.5 Å
2x 3D classification
Mg2+ lower,118,618 particles
GSFSC Resolution: 3.15Å
4.0 Å
3.2 Å
2.7 Å
Orientation
distribution plot
Local refinement
with TMD mask
Mg2+ upper, 105,462 particles
GSFSC Resolution: 3.21Å
4.0 Å
3.2 Å
2.7 Å
Orientation
distribution plot
e
Mg2+: NU refinement, non-active state
V656
I655
V656
I655
F627
T626
S626
T626
GluN1a
Chain A
626-656
GluN2B
Chain A
625-655
GluN1a
Chain A
626-656
GluN2B
Chain A
625-655
f
Mg2+ lower: local refinement, TMD
V656
I655
V656
I655
F627
T626
S626
T626
GluN1a
Chain A
626-656
GluN2B
Chain A
625-655
GluN1a
Chain A
626-656
GluN2B
Chain A
625-655
g
Mg2+ upper: local refinement, TMD
V656
I655
V656
I655
F627
T626
S626
T626
GluN1a
Chain A
626-656
GluN2B
Chain A
625-655
GluN1a
Chain A
626-656
GluN2B
Chain A
625-655

**Extended Data Figure 2. Single particle cryo-EM of divalent cation-free EDTA-treated, and the $Mg^{2+}$-bound GluN1a-2B NMDAR (Related to Figure 1 and Figure 2). a,** Single-particle cryo-EM workflow of the divalent cation-free EDTA-treated GluN1a-2B NMDAR. **b,** Map quality assessment of the divalent cation-free EDTA-treated GluN1a-2B NMDAR NU-refinement at the M3/M3' region. **c,** Map quality assessment of the divalent cation-free EDTA-treated GluN1a-2B NMDAR TMD local refinement at the M3/M3' region. **d,** Single-particle cryo-EM workflow of the $Mg^{2+}$-bound GluN1a-2B NMDAR. To obtain $Mg^{2+}$-bound GluN1a-2B NMDAR, two rounds of 3D classification without alignment were performed to obtain homogenous $Mg^{2+}$-bound states. Each class was further processed with TMD-masked local refinement for improved resolution. **e,** Map quality assessment of the $Mg^{2+}$-bound GluN1a-2B NMDAR NU-refinement at the M3/M3' region. **f-g,** Map quality assessment of the $Mg^{2+}$-bound GluN1a-2B NMDAR TMD local refinement at the M3/M3' region. The scale bars in the micrographs (panel a and d) represent 20 nm.

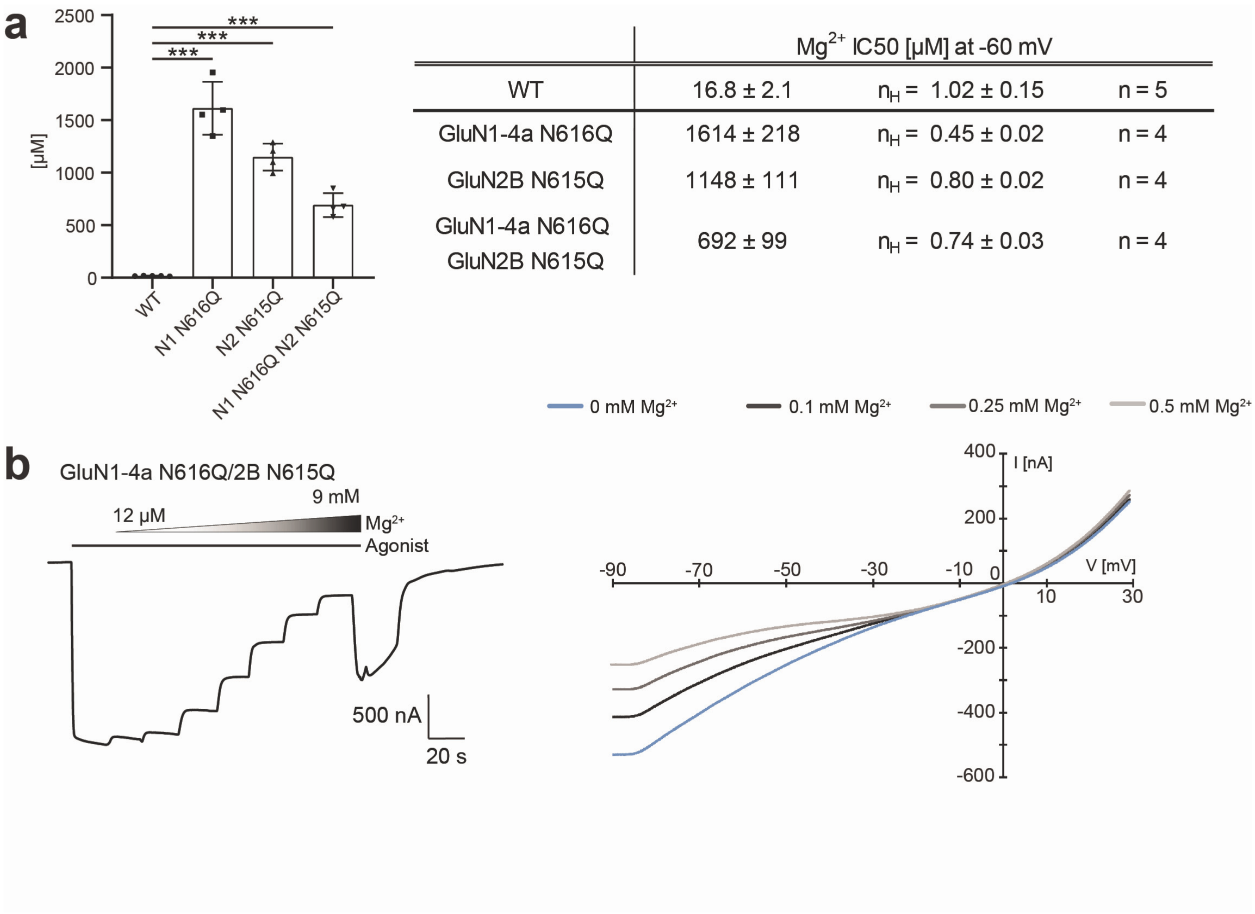


| | Mg$^{2+}$ IC50 [µM] at -60 mV | | |
|---|---|---|---|
| WT | 16.8 ± 2.1 | $n_H$ = 1.02 ± 0.15 | n = 5 |
| GluN1-4a N616Q | 1614 ± 218 | $n_H$ = 0.45 ± 0.02 | n = 4 |
| GluN2B N615Q | 1148 ± 111 | $n_H$ = 0.80 ± 0.02 | n = 4 |
| GluN1-4a N616Q GluN2B N615Q | 692 ± 99 | $n_H$ = 0.74 ± 0.03 | n = 4 |

**Extended Data Fig. 3. Effects of the Asn-cage mutations on voltage-dependent Mg$^{2+}$ block (Related to Figure 2). a,** $IC_{50}$ values ± SD derived from Mg$^{2+}$ concentration-response curves at -60 mV through TEVC. The statistical analysis was performed by One-Way ANOVA (*** $p < 0.001$, ** $0.001 < p < 0.01$, * $0.01 < p < 0.05$, n.s. not significant). The table lists the $IC_{50}$ values and Hill coefficients ± SD ($n_H$) calculated based on the dose-response curves. $IC_{50}$ values were calculated from independent dose-response recordings from at least four independent oocytes (n). **b,** Whole-cell TEVC electrophysiology on cRNA-injected *Xenopus laevis* oocytes expressing GluN1a Asn616Gln-2B Asn615Gln NMDAR at -60 mV and I/V recording at -90 - +30 mV (ramp = 2 s). The isotype of GluN1a used in these TEVC experiments is GluN1-4a.

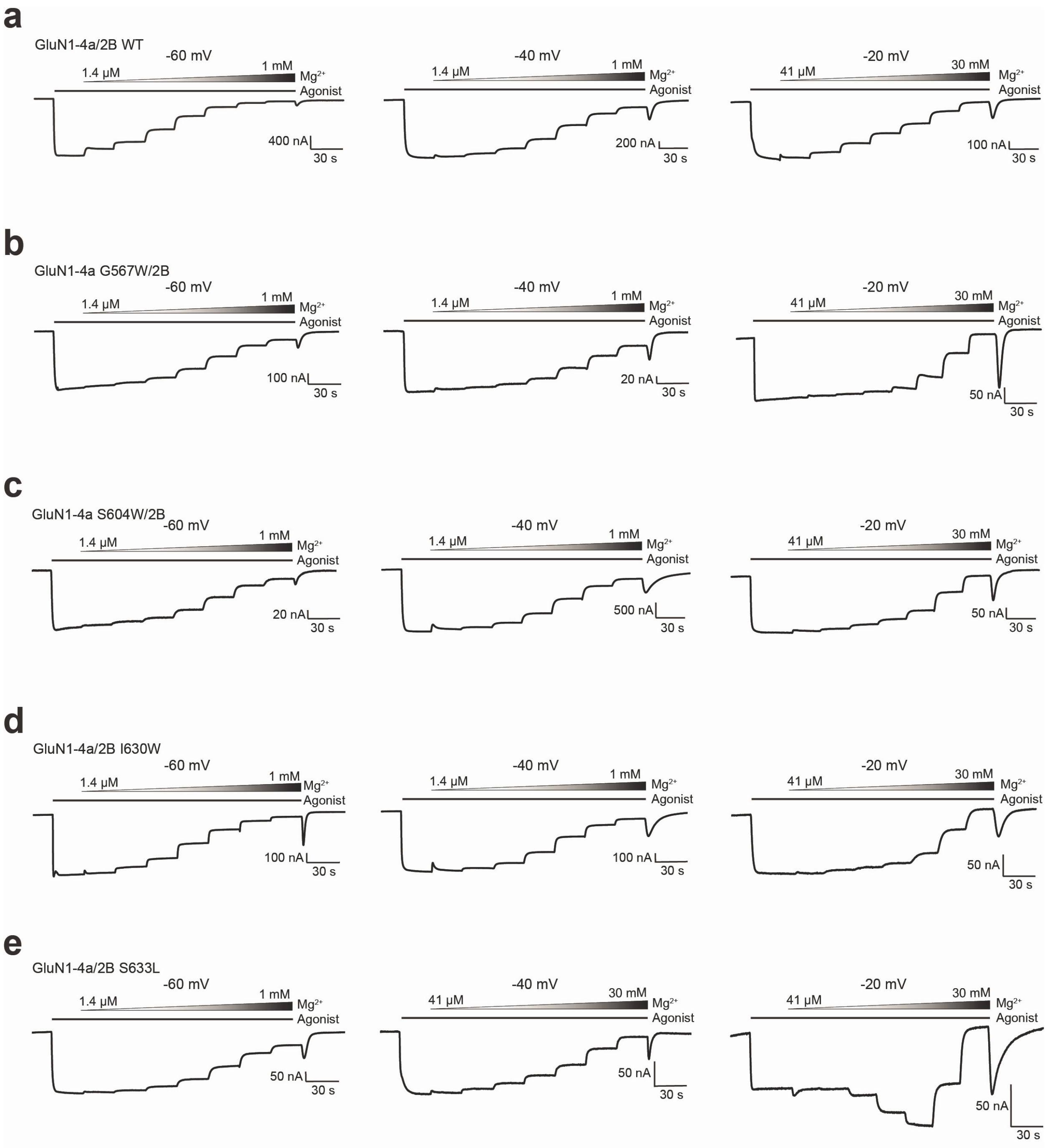


**Extended Data Figure 4. $IC_{50}$ evaluation of the $Mg^{2+}$-block on GluN1a-2B NMDAR (related to Figure 4).** **a-e,** $IC_{50}$ evaluation through TEVC electrophysiology on cRNA-injected (**a**: GluN1a-2B NMDAR wild-type, **b**: GluN1a Gly567Trp/2B NMDAR, **c**: GluN1a Ser604Trp/2B NMDAR, **d**: GluN1a/2B Ile630Trp NMDAR, **Ee**: GluN1a/2B Ser633Leu NMDAR) oocytes at -60, -40, and -20 mV. The isotype of GluN1a used in these TEVC experiments is GluN1-4a.

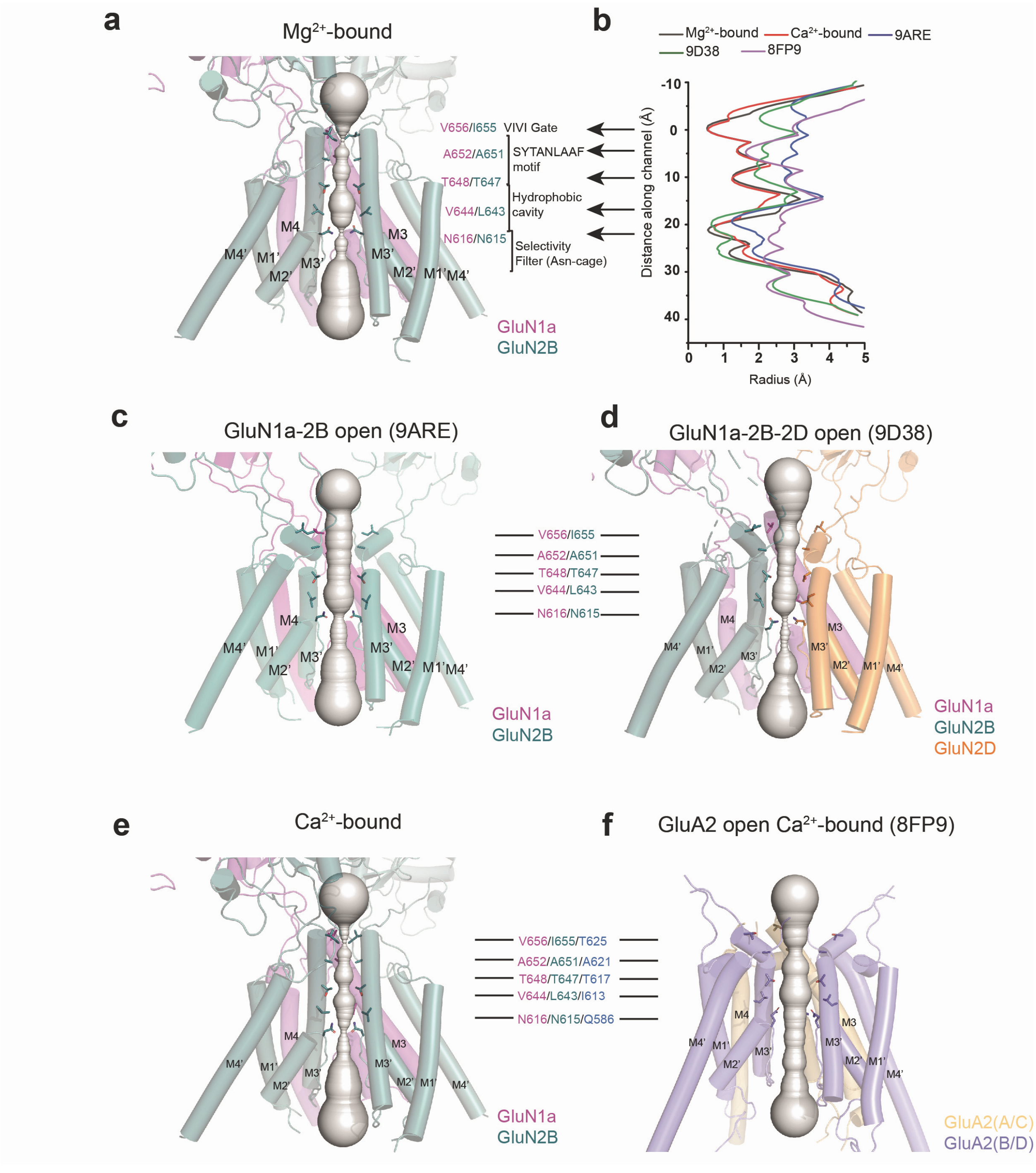


**Extended Data Fig. 5. Comparison of iGluR pores. a-b,** Hole analysis of the GluN1a-2B NMDAR bound to $Mg^{2+}$ and agonists (a) and pore-radius measurement (b). The TMD motifs are annotated. **c-f**, Hole analysis of GluN1a-2B open channel (PDB code: 9ARE, panel c), GluN1a-2B-2D open channel (PDB code: 9D38, panel d), $Ca^{2+}$-bound GluN1a-2B (panel d), and GluA2 open channel bound to $Ca^{2+}$ (PDB code: 8FP9, panel f).

**Extended Data Table 1. Cryo-EM data collection and statistics**

| | EDTA buffered GluN1a-2B NMDAR (EMDB: XXX) (PDB: XXX) | EDTA buffered GluN1a-2B NMDAR TMD (EMDB: XXX) (PDB: XXX) | $Ca^{2+}$ conditioned GluN1a-2B NMDAR (EMDB: XXX) (PDB: XXX) | $Ca^{2+}$ State-1 GluN1a-2B NMDAR TMD (EMDB: XXX) (PDB: XXX) | $Ca^{2+}$ State-2 GluN1a-2B NMDAR TMD (EMDB: XXX) (PDB: XXX) | $Ca^{2+}$ State-3 GluN1a-2B NMDAR TMD (EMDB: XXX) (PDB: XXX) |
|---|---|---|---|---|---|---|
| **Data collection and processing** | | | | | | |
| Microscope | Titan Krios | Titan Krios | Titan Krios | Titan Krios | Titan Krios | Titan Krios |
| Camera | K3/CDS | K3/CDS | K3/CDS | K3/CDS | K3/CDS | K3/CDS |
| Magnification | 105K | 105K | 105K | 105K | 105K | 105K |
| Energy filter slit width (eV) | 14 | 14 | 14 | 14 | 14 | 14 |
| Collection software | EPU | EPU | EPU | EPU | EPU | EPU |
| Voltage (kV) | 300 | 300 | 300 | 300 | 300 | 300 |
| Cumulative exposure ($e^-$/Å$^2$) | 58.4 | 58.4 | 55.6-64.0 | 58.5 | 55.6-64.0 | 55.6-64.0 |
| Exposure rate ($e^-$/Å$^2$/frame) | 1.46 | 1.46 | 1.39-1.6 | 1.95 | 1.39-1.6 | 1.39-1.6 |
| Defocus range (μm) | -2.2~ -0.6 | -2.2~ -0.6 | -2.2~ -0.6 | -2.2~ -0.6 | -2.2 ~ -0.6 | -2.2 ~ -0.6 |
| Pixel size (Å) | 0.827 | 0.827 | 0.84 | 0.827 | 0.84 | 0.84 |
| Symmetry imposed | C1 | C2 | C1 | C1 | C2 | C2 |
| Number of micrographs | 12,035 | 12,035 | 30,928 | 18,275 | 30,928 | 30,928 |
| Initial particle images (no.) | 2,826,035 | 2,826,035 | 7,062,525 | 4,017,866 | 7,062,525 | 7,062,525 |
| Final particle images (no.) | 131,512 | 131,512 | 823,417 | 55,995 | 116,195 | 61,249 |
| 0.143 FSC map masked (Å) | 3.18 | 3.01 | 2.59 | 3.60 | 2.81 | 2.97 |
| 0.143 FSC map unmasked(Å) | 3.8 | 4.0 | 3.1 | 5.9 | 3.9 | 4.2 |
| **Refinement** | | | | | | |
| Refinement package | Phenix | Phenix | Phenix | Phenix | Phenix | Phenix |
| Initial model used (PDB code) | 7SAA | 7SAA | 7SAA | 7SAA | 7SAA | 7SAA |
| Map sharpening B factor (Å$^2$) | -99.7 | -107.1 | -91.2 | -105.4 | -89.0 | -98.0 |
| Model composition | | | | | | |
| Non-hydrogen atoms | 20853 | 3,784 | 21,064 | 3,464 | 3,997 | 3,751 |
| Protein residues | 3157 | 518 | 3,166 | 516 | 527 | 518 |
| Ligands | 1 | 2 | 0 | 1 | 3 | 1 |
| Water | 0 | 28 | 0 | 9 | 37 | 23 |
| CC map vs. model | 0.85 | 0.88 | 0.76 | 0.76 | 0.85 | 0.81 |
| R.m.s. deviations | | | | | | |
| Bond lengths (Å) | 0.005 | 0.003 | 0.001 | 0.004 | 0.002 | 0.004 |
| Bond angles (°) | 0.498 | 0.415 | 0.371 | 0.584 | 0.422 | 0.576 |
| Validation | | | | | | |
| MolProbity score | 1.82 | 1.39 | 1.22 | 2.00 | 1.11 | 1.60 |
| Clashscore | 6.87 | 3.80 | 2.48 | 10.52 | 3.16 | 7.78 |
| Rotamer outliers (%) | 0.78 | 0.00 | 0.56 | 0.74 | 0.00 | 0.56 |
| Ramachandran plot | | | | | | |
| Favored (%) | 93.23 | 96.56 | 96.88 | 92.68 | 98.01 | 96.96 |
| Allowed (%) | 6.58 | 3.04 | 2.93 | 6.71 | 1.99 | 3.04 |
| Outliers (%) | 0.19 | 0.40 | 0.19 | 0.61 | 0.00 | 0.00 |
| CaBLAM outliers (%) | 3.25 | 1.91 | 2.79 | 3.85 | 1.46 | 1.91 |

### Extended Data Table 1. Cryo-EM data collection and statistics

| | $Ca^{2+}$ State-4 GluN1a-2B NMDAR TMD (EMDB: XXX) (PDB: XXX) | $Ca^{2+}$ State-5 GluN1a-2B NMDAR TMD (EMDB: XXX) (PDB: XXX) | $Mg^{2+}$ conditioned GluN1a-2B NMDAR (EMDB: XXX) (PDB: XXX) | $Mg^{2+}$ lower GluN1a-2B NMDAR TMD (EMDB: XXX) (PDB: XXX) | $Mg^{2+}$ upper GluN1a-2B NMDAR TMD (EMDB: XXX) (PDB: XXX) |
|---|---|---|---|---|---|
| **Data collection and processing** | | | | | |
| Microscope | Titan Krios | Titan Krios | Titan Krios | Titan Krios | Titan Krios |
| Camera | K3/CDS | K3/CDS | K3/CDS | K3/CDS | K3/CDS |
| Magnification | 105K | 105K | 105K | 105K | 105K |
| Energy filter slit width (eV) | 14 | 14 | 14-20 | 14-20 | 14-20 |
| Collection software | EPU | EPU | EPU | EPU | EPU |
| Voltage (kV) | 300 | 300 | 300 | 300 | 300 |
| Cumulative exposure ($e^-$/Å$^2$) | 55.6-64.0 | 55.6-64.0 | 58.2-71.7 | 58.2-71.7 | 58.2-71.7 |
| Exposure rate ($e^-$/Å$^2$/frame) | 1.39-1.6 | 1.39-1.6 | 1.94-2.39 | 1.94-2.39 | 1.94-2.39 |
| Defocus range (µm) | -2.2~ -0.6 | -2.2~ -0.6 | -2.6~ -0.8 | -2.6~ -0.8 | -2.6 ~ -0.8 |
| Pixel size (Å) | 0.84 | 0.84 | 0.856 | 0.856 | 0.856 |
| Symmetry imposed | C2 | C2 | C1 | C2 | C2 |
| Number of micrographs | 30,928 | 30,928 | 38,717 | 38,717 | 38,717 |
| Initial particle images (no.) | 7,062,525 | 7,062,525 | 5,106,550 | 5,106,550 | 5,106,550 |
| Final particle images (no.) | 102,122 | 87,601 | 512,800 | 118,618 | 105,462 |
| 0.143 FSC map masked (Å) | 2.76 | 2.69 | 3.16 | 3.15 | 3.21 |
| 0.143 FSC map unmasked(Å) | 4.0 | 4.0 | 3.6 | 4.1 | 4.1 |
| **Refinement** | | | | | |
| Refinement package | Phenix | Phenix | Phenix | Phenix | Phenix |
| Initial model used (PDB code) | 7SAA | 7SAA | 7SAA | 7SAA | 7SAA |
| Map sharpening B factor (Å$^2$) | -78.6 | -88.0 | -118.8 | -127.0 | -115.4 |
| Model composition | | | | | |
| Non-hydrogen atoms | 3,791 | 3726 | 21219 | 3,791 | 3,666 |
| Protein residues | 518 | 516 | 3161 | 508 | 514 |
| Ligands | 1 | 1 | 0 | 1 | 1 |
| Water | 32 | 25 | 0 | 14 | 23 |
| CC map vs. model | 0.86 | 0.86 | 0.78 | 0.82 | 0.84 |
| R.m.s. deviations | | | | | |
| Bond lengths (Å) | 0.002 | 0.002 | 0.004 | 0.002 | 0.003 |
| Bond angles (°) | 0.438 | 0.465 | 0.496 | 0.422 | 0.518 |
| Validation | | | | | |
| MolProbity score | 1.43 | 1.50 | 1.49 | 0.81 | 1.25 |
| Clashscore | 5.26 | 7.40 | 3.54 | 1.09 | 1.97 |
| Rotamer outliers (%) | 0.27 | 0.85 | 0.36 | 0.31 | 0.60 |
| Ramachandran plot | | | | | |
| Favored (%) | 97.17 | 97.56 | 94.96 | 98.35 | 95.92 |
| Allowed (%) | 2.83 | 2.44 | 4.97 | 1.45 | 4.08 |
| Outliers (%) | 0.00 | 0.00 | 0.06 | 0.21 | 0.00 |
| CaBLAM outliers (%) | 1.28 | 1.92 | 3.50 | 0.87 | 2.15 |

**Extended Data Table 2. $IC_{50}$ [μM] values of lipid-binding site mutants**

| [mV] | -60 | -40 | -20 |
|---|---|---|---|
| WT | 16.8 ± 2.1<br>$n_H$ = 0.94 ± 0.03 ; n = 5 | 79.1 ± 4.5<br>$n_H$ = 0.88 ± 0.02 ; n = 5 | 1662 ± 52<br>$n_H$ = 0.78 ± 0.06 ; n = 6 |
| GluN1-4a V566W | 44.9 ± 4.4<br>$n_H$ = 0.81 ± 0.10 ; n = 6 | 243.9 ± 11.2<br>$n_H$ = 0.75 ± 0.05 ; n = 5 | 1042 ± 236<br>$n_H$ = 1.13 ± 0.23 ; n = 4 |
| GluN1-4a G567W | 35.0 ± 4.3<br>$n_H$ = 0.84 ± 0.06 ; n = 5 | 201.8 ± 18.8<br>$n_H$ = 0.84 ± 0.13 ; n = 4 | 4989 ± 488<br>$n_H$ = 1.62 ± 0.10 ; n = 5 |
| GluN1-4a S604W | 20.9 ± 1.6<br>$n_H$ = 0.93 ± 0.02 ; n = 5 | 91.9 ± 4.0<br>$n_H$ = 0.88 ± 0.02 ; n = 5 | 5915 ± 342<br>$n_H$ = 1.14 ± 0.03 ; n = 4 |
| GluN1-4a M607W | 41.6 ± 1.2<br>$n_H$ = 0.93 ± 0.04 ; n = 5 | 174.2 ± 12.6<br>$n_H$ = 0.80 ± 0.02 ; n = 5 | 7041 ± 677<br>$n_H$ = 1.77 ± 0.12 ; n = 5 |
| GluN1-4a L615W | 24.4 ± 1.9<br>$n_H$ = 0.99 ± 0.05 ; n = 4 | 105.3 ± 3.5<br>$n_H$ = 0.94 ± 0.04 ; n = 4 | 1018 ± 146<br>$n_H$ = 0.95 ± 0.11 ; n = 6 |
| GluN1-4a L615Q | 48.7 ± 3.8<br>$n_H$ = 0.97 ± 0.03 ; n = 6 | 264.6 ± 23.8<br>$n_H$ = 0.91 ± 0.06 ; n = 4 | 2365 ± 370<br>$n_H$ = 1.48 ± 0.13 ; n = 4 |
| GluN2B T626W | 14 ± 1.9<br>$n_H$ = 0.93 ± 0.06 ; n = 4 | 89.7 ± 5.0<br>$n_H$ = 0.85 ± 0.02 ; n = 5 | 2176 ± 490<br>$n_H$ = 1.04 ± 0.04 ; n = 5 |
| GluN2B I630W | 23.4 ± 1.7<br>$n_H$ = 1.02 ± 0.07 ; n = 5 | 79.3 ± 2.9<br>$n_H$ = 0.92 ± 0.02 ; n = 4 | 5655 ± 476<br>$n_H$ = 1.68 ± 0.06 ; n = 4 |
| GluN2B S633L | 164.3 ± 11.4<br>$n_H$ = 0.80 ± 0.04 ; n = 5 | 3015 ± 184<br>$n_H$ = 0.82 ± 0.03 ; n = 4 | 10472 ± 1068<br>$n_H$ = n.d. ; n = 5 |
| GluN2B F637W | 26.9 ± 2.3<br>$n_H$ = 1.03 ± 0.04 ; n = 4 | 95.6 ± 10.2<br>$n_H$ = 1.10 ± 0.06 ; n = 5 | 661.7 ± 29.4<br>$n_H$ = 1.57 ± 0.06 ; n = 4 |

$IC_{50}$ values ± SD derived from $Mg^{2+}$ concentration-response curves at -60, -40, -20 mV through TEVC. The table lists the $IC_{50}$ values and Hill coefficients ± SD ($n_H$) calculated based on the dose-response curves. $IC_{50}$ values were calculated from independent dose-response recordings from at least four independent oocytes (n). The isotype of GluN1a used in these TEVC experiments is GluN1-4a.
